# Necator americanus Ancylostoma secreted protein-2 (Na-ASP-2) selectively binds an ascaroside (ascr#3)

**DOI:** 10.1101/2020.08.07.224964

**Authors:** Ola El Atab, Rabih Darwiche, Nathanyal J. Truax, Roger Schneiter, Kenneth G. Hull, Daniel Romo, Oluwatoyin A. Asojo

**Affiliations:** Division of Biochemistry, Department of Biology, University of Fribourg, Chemin du Musée 10, CH 1700 Fribourg, Switzerland; Department of Biological Chemistry and Molecular Pharmacology, Harvard Medical School, Boston, MA 02115; Department of Chemistry and Biochemistry & The CPRIT Synthesis and Drug-Lead Discovery Laboratory, Baylor University, One Bear Place, Waco, Texas 76798-7348, United States; Department of Chemistry and Biochemistry, 200 William Harvey Way, Hampton University, Hampton VA, 23668; National School of Tropical Medicine, Baylor College of Medicine, Houston Texas, 77030

**Keywords:** Venom allergen-like (VAL), sperm coating protein (SCP), TAPs [testis specific proteins (Tpx) / antigen 5 (Ag5) / pathogenesis related-1 (PR-1) / Sc7], CAP [<u>c</u>ysteine-rich secretory protein (CRISP) / <u>a</u>ntigen 5 / <u>p</u>athogenesis related-1 (PR-1)], lipid binding, SCP/TAPS (Sperm-coating protein / Tpx / antigen 5 / pathogenesis related-1 / Sc7), ascarosides

## Abstract

During their infective stages, hookworms release excretory-secretory (E-S) products, including small molecules and proteins, to help evade and suppress the host’s immune system. Small molecules found in E-S products of mammalian hookworms include nematode derived metabolites like ascarosides, which are composed of the sugar ascarylose linked to a fatty acid side chain. Ascarosides play vital roles in signaling, development, reproduction, and survival. The most abundant proteins found in hookworm E-S products are members of the protein family known as *Ancylostoma* secreted protein (ASP). ASP belongs to the SCP/TAPS (sperm-coating protein / Tpx / antigen 5 / pathogenesis related-1 / Sc7) superfamily of proteins, members of which have previously been shown to bind to eicosanoids and fatty acids. These molecules are structurally similar to the fatty acid moieties of ascarosides. The objective of this study was to determine if the hookworm ASP; *N. americanus Ancylostoma* secreted protein 2 (*Na*-ASP-2) binds to the ascarosides or their fatty acid moieties. We describe investigations of our hypothesis that there is a functional relationship between the major secreted proteins and signaling small molecules found in hookworm E-S products. To accomplish this, several ascarosides and their fatty acid moieties were synthesized and tested for *in vitro* binding to *Na*-ASP-2 using a ligand competition assay and microscale thermophoresis. Our results reveal that the fatty acid moieties of the ascarosides, bind specifically to the palmitic acid binding cavity of *Na*-ASP-2. Additionally, ascr#3, an ascaroside that is present in mammalian hookworm E-S products binds to the palmitic acid binding cavity of *Na*-ASP-2, whereas oscr#10 which is not found in hookworm E-S products does not bind. Future studies are required to determine the structural basis of ascaroside binding by *Na*-ASP-2 and to understand the physiological significance of these observations.

## 1. Introduction

*Necator americanus* and *Ancylostoma duodenale* are hookworms that infect more than 400 million of the world’s poorest people causing a disease burden of over 22 million disability-adjusted life years (de Silva et al., 2003; Diemert et al., 2018; Hotez, 2007; Murray et al., 2014). During the transition to parasitism, the most abundant proteins secreted by third-stage infective larvae (L3) of *N. americanus* upon host entry are *N. americanus Ancylostoma* secreted protein 1 (*Na*-ASP-1) and *N. americanus Ancylostoma* secreted protein 2 (*Na*-ASP-2) (Hotez et al., 2003). These *Ancylostoma* secreted protein sometimes referred to as VALs (venom allergen like) are the major protein components of the L3 excretory-secretory (E-S) products that facilitate the evasion and suppression of the host’s immune system and have been found in all parasitic nematodes studied to date (Asojo et al., 2018; Darwiche et al., 2018; Gao et al., 2001; Hawdon and Hotez, 1996; Hawdon et al., 1996; Hawdon et al., 1995; Hawdon et al., 1999; Zhan et al., 2003). ASPs belong to the SCP/TAPS (sperm-coating protein / Tpx / antigen 5 / pathogenesis related-1 / Sc7) superfamily of proteins, NCBI domain cd00168 or Pfam PF00188 (Gibbs et al., 2008). SCP/TAPS proteins include plant PR-1 (pathogenesis-related 1) and CRISPs (cysteine-rich secretory protein), which are expressed in the mammalian reproductive tract, and venom allergens from insects and reptiles (Gibbs and O’Bryan, 2007; Gibbs et al., 2008; Gibbs et al., 2006). Members of the SCP/TAPS superfamily are also implicated in other biological phenomena including cellular defense such as plant responses to pathogens, sexual reproduction, and human brain tumor growth (Ding et al., 2000; Gao et al., 2001; Gibbs et al., 2010; Gibbs et al., 2008; Hawdon et al., 1999; Zhan et al., 2003).

SCP/TAPS proteins have either one or two ∼15 kDa cysteine-rich CAP domains (cysteine-rich secretory protein, antigen 5, and pathogenesis-related 1) as typified by the structures of *Na*-ASP-2 (one CAP domain) and *Na*-ASP-1 (two covalently linked CAP domains) (Asojo, 2011; Asojo et al., 2005a; Asojo et al., 2011; Borloo et al., 2013; Fernandez et al., 1997; Gibbs et al., 2008; Guo et al., 2005; Serrano et al., 2004; Shikamoto et al., 2005; Wang et al., 2005; Xu et al., 2012). The CAP domain has an alpha-beta-alpha sandwich topology, and up to 50 % loop regions, which often makes it difficult to predict structure by homology modelling alone (Asojo et al., 2005a; Asojo et al., 2005b; Darwiche et al., 2016; Kelleher et al., 2014). The CAP domain has multiple cavities and verified ligand binding regions, and the first to be identified was a large central cavity that may contain a tetrad of residues, two His and two Glu that bind divalent cations including Zn^2+^ and Mg^2+^(Asojo et al., 2018; Asojo et al., 2011; Darwiche et al., 2018; Gibbs et al., 2008; Mason et al., 2014; Wang et al., 2010). Distinct lipid binding sites have been verified in SCP/TAPS proteins, including phosphatidylinositol binding regions on the surface of human golgi-associated plant pathogenesis-related protein 1 (GAPR-1) (Darwiche et al., 2016; van Galen et al., 2012; Van Galen et al., 2010; Xu et al., 2012). A sterol binding caveolin-binding motif (CBM) of the yeast CAP proteins required for *in vivo* transport of cholesterol has also been identified in diverse SCP/TAPS proteins (Asojo et al., 2018; Choudhary et al., 2014; Darwiche et al., 2017a; Darwiche et al., 2016; Darwiche et al., 2018; Darwiche et al., 2017b; Kelleher et al., 2014). Furthermore, a hydrophobic channel that binds leukotrienes with sub-micromolar affinities, allows tablysin-15, an SCP/TAPS protein from the horsefly *Tabanus yao*, to function as an anti-inflammatory scavenger of eicosanoids (Xu et al., 2012). This binding cavity is formed by conserved central helices in SCP/TAPS proteins and also binds fatty acids, including palmitic acid (Xu et al., 2012). Our previous studies revealed that palmitic acid specifically binds to this cavity in other SCP/TAPS proteins including pathogen-related in yeast protein 1(Pry1) from *Saccharomyces cerevisiae* hence the cavity is referred to as the fatty acid-binding cavity (Asojo et al., 2018; Darwiche et al., 2016; Darwiche et al., 2018; Kelleher et al., 2014). While lipid binding had been confirmed for other parasite SCP/TAPS proteins, we solved the structure of *Na*-ASP-2 prior to the discovery of the lipid binding activity of members of this protein superfamily (Asojo et al., 2005a). The impetus for the current studies is to unravel possible lipid binding functions of *Na*-ASP-2. Our working hypothesis is that there is a functional relationship between small molecules and proteins secreted in hookworm E-S products. Thus, we are interested in the ability of *Na*-ASP-2 to bind small molecules with known functions secreted by L3 hookworms. These small molecules include nematode derived metabolites notably the ascarosides which regulate a diverse range of phenotypes in nematodes including dauer arrest, mate attraction, aggregation and olfactory plasticity (Butcher et al., 2007; Choe et al., 2012; Gallo and Riddle, 2009; Hollister et al., 2013; Izrayelit et al., 2012; Jezyk and Fairbairn, 1967; Kaplan et al., 2011; Kunert, 1992; Ludewig and Schroeder, 2013; Noguez et al., 2012; Rhoads et al., 2015; Sakai et al., 2013; Tarr and Fairbairn, 1973; Tarr and Schnoes, 1973). Ascarosides are multifunctional small molecules that interact with G-protein–coupled receptors (GPCRs) (Butcher, 2017; Park et al., 2012). Ascarosides are composed of the sugar ascarylose linked to a fatty acid moiety (eg. ascr#3 (**1**) and oscr#10 (**2**), Figure 1) and while ascr#3 (**1**) was identified in the E-S products of mammalian hookworms, oscr#10 (**2**) was identified in other nematodes, but not hookworm (Choe et al., 2012; Gallo and Riddle, 2009; Hollister et al., 2013; Izrayelit et al., 2012; Jezyk and Fairbairn, 1967; Kaplan et al., 2011; Kunert, 1992; Ludewig and Schroeder, 2013; Noguez et al., 2012; Rhoads et al., 2015; Tarr and Fairbairn, 1973; Tarr and Schnoes, 1973). Since the fatty acid moieties of ascarosides are similar to those that are capable of binding to the fatty acid-binding cavity of ASPs, we carried out studies to determine if the ascarosides or their fatty acid moieties bind to *Na*-ASP-2.

**Figure 1.**
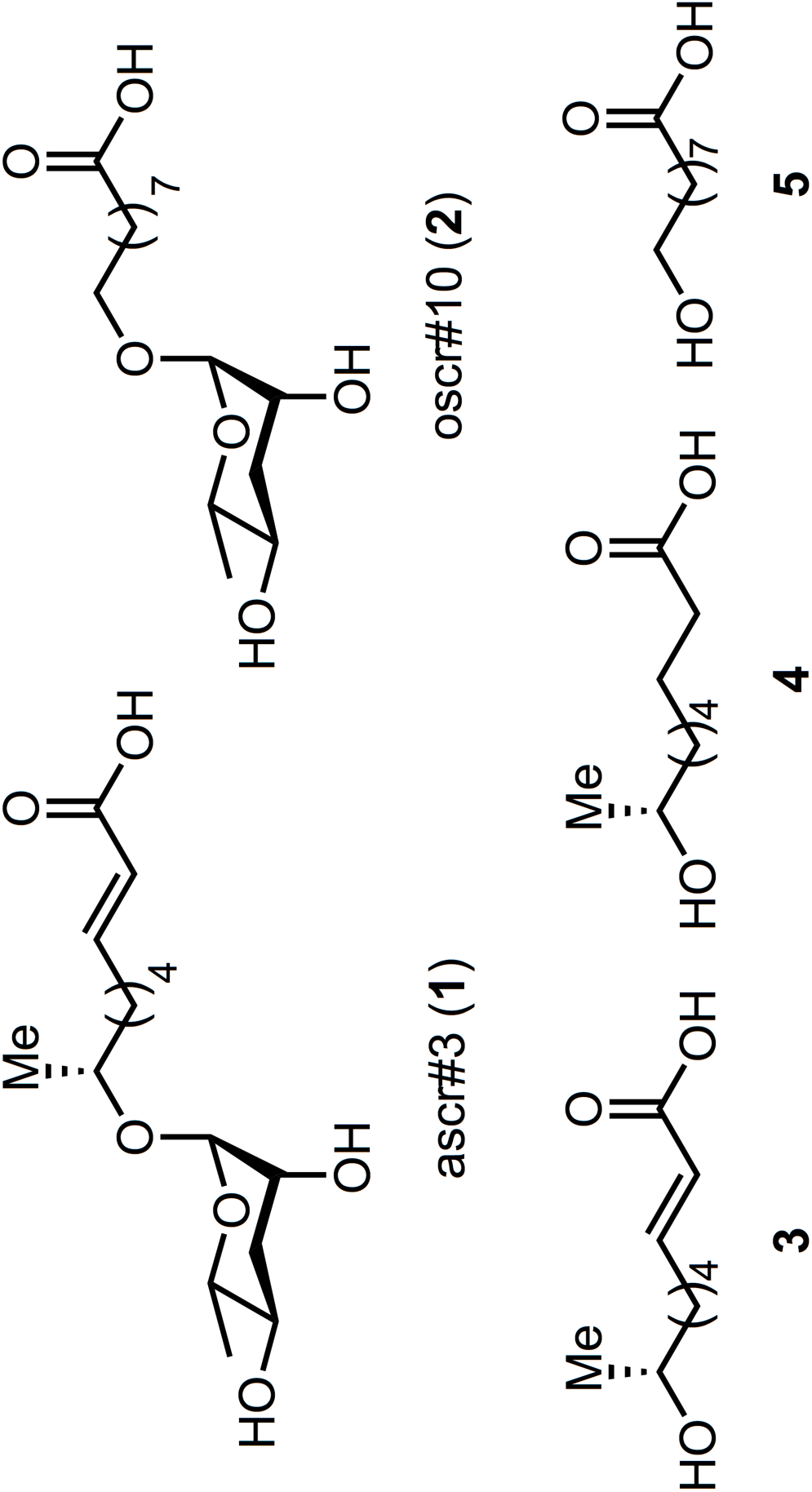
Targeted ascarosides and their fatty acid side moieties. The corresponding ascarosides are ascr#3 (**1**); oscr#10 (**2**) and their side chain moieties are **3-5**. Compound names are **3** = (R)-8-hydroxynonanoic acid, **4** = (R,E)-8-hydroxynon-2-enoic acid, and **5** = 9-hydroxynonanoic acid.

## 2. Experimental Procedures

### 2.1. Expression and purification of Pry1 and *Na*-ASP-2

DNA encoding for Pry1 and *Na*-ASP-2 were PCR amplified and cloned into NcoI and XhoI restriction sites of pET22b vector (Novagen, Merck, Darmstadt, Germany), which contains a pelB signal sequence to direct the secretion of expressed protein into the periplasmic space. Plasmids were transformed into *Escherichia coli* BL21 and proteins were expressed with a C-terminal polyhistidine-tag. Protein expression was induced overnight with lactose at 24°C. Cells were collected, lysed and incubated with nickel-nitrilotriacetic acid beads as per the manufacturer instructions (Qiagen, Hilden, Germany). Beads were washed, loaded onto a Ni^2+^-NTA column and proteins were eluted in 60 mM NaH_2_PO_4_, 300 mM NaCl and 300 mM imidazole, pH 8.0. Prior to microscale thermophoresis experiments, proteins were applied to Zeba^™^ spin desalting columns (Thermo scientific) and the buffer was exchanged to 60 mM NaH_2_PO_4_, 300 mM NaCl, pH 8.0. Protein concentration was determined by Lowry assay using folin reagent and BSA as standard.

### 2.2. *In vitro* radioligand lipid binding assay

The radioligand binding assay was performed as described previously (Choudhary and Schneiter, 2012; Im et al., 2005). 100 pmol of purified untagged CAP protein (*Na*-ASP-2 or Pry1) in binding buffer (20 mM Tris, pH 7.5, 30 mM NaCl, 0.05% Triton X-100) was incubated for 1 h at 30 °C with different concentrations of either [^3^H]-cholesterol or [^3^H]-palmitic acid. Protein was removed from unbound ligand by adsorption to Q-sepharose beads (GE healthcare, USA), the beads were washed, protein was eluted and the protein-bound radioligand was quantified by scintillation counting. For competition binding assays, specified concentrations of unlabeled cholesterol, palmitic acid or ligands, were included in the binding reaction. Non-specific binding was determined by performing the assays without the addition of protein. Statistical significance of data was analyzed by multiple t-test (GraphPad Prism, La Jolla, CA).

### 2.3. Microscale Thermophoresis

Microscale thermophoresis was performed using a Monolith NT.115 from Nanotemper Technologies (Munich, Germany) (Seidel et al., 2012; Shang et al., 2012; Zillner et al., 2012). His-tagged protein (Pry1 or *Na*-ASP-2) was fluorescently labeled using the RED-tris-NTA His tag protein labeling kit (Nanotemper Technologies). Labeled protein (Pry1 or *Na*-ASP-2) was subsequently added to serial dilution of unlabeled ligand (ascarosides or their fatty acid moieties) in binding buffer (20 mM Tris pH 7.5, 30 mM NaCl, 0.05% Triton X-100). Each sample was loaded into standard glass capillaries, and measurements were performed at 60% power setting. The dissociation constant Kd was obtained by plotting the normalized fluorescence (Fnorm) against the logarithm of ligand concentration. Experiments were performed in triplicates and data were fitted using the Kd model with the MO.Affinity Analysis software (Nanotemper Technologies, Munich, Germany).

### 2.4. Synthesis of Ascarosides and ligands

Benzoyl protected ascarylose **8** was prepared as previously reported by Jeong and co-workers from commercially available L-rhamnose **6** (Jeong et al., 2005) with the exception of a modified final reduction (Figure 2). The previously reported reduction of lactone **7** with disiamyl borane (Jeong et al., 2005) proved irreproducible in our hands, resulting in incomplete conversion and low overall yields (∼40 %). Thus, an alternative was identified involving reduction with 9-BBN to provide the desired lactol **7** in improved yield (70 %). With protected ascarylose **8** in hand, we next studied glycosylation at C1 to append the fatty acid side chain present in the targeted ascarosides. Previous synthetic strategies to these targets involved glycosylation of secondary alcohols bearing long alkyl chains with a terminal alkene which was subsequently utilized for late stage cross metathesis or oxidations (Butcher et al., 2009; Hollister et al., 2013; Jeong et al., 2005; Martin et al., 2009; Noguez et al., 2012; Srinivasan et al., 2012). Since we intended to study the binding affinity of the natural ascarocides and their intact fatty acid moieties independently, we decided to first synthesize intact fatty acid side moieties **9** and **11** and then couple them directly to protected ascarylose **8** during the penultimate step of the sequence. This strategy provided rapid access to ascarosides **1** and **2** along with fatty acid derivatives **3-5** for screening. Subsequent Lewis acid-mediated glycosylation with BF_3_•Et_2_O of fatty acid **9** (see SI for synthetic details) and commercially available acid **11** (Jeong et al., 2005) proceeded uneventfully and provided protected ascarosides **10** and **12** in 68 and 66% yield, respectively. Subsequent global deprotection with lithium hydroxide gave ascr#3 (**1**) and oscr#10 (**2**).

**Figure 2.**
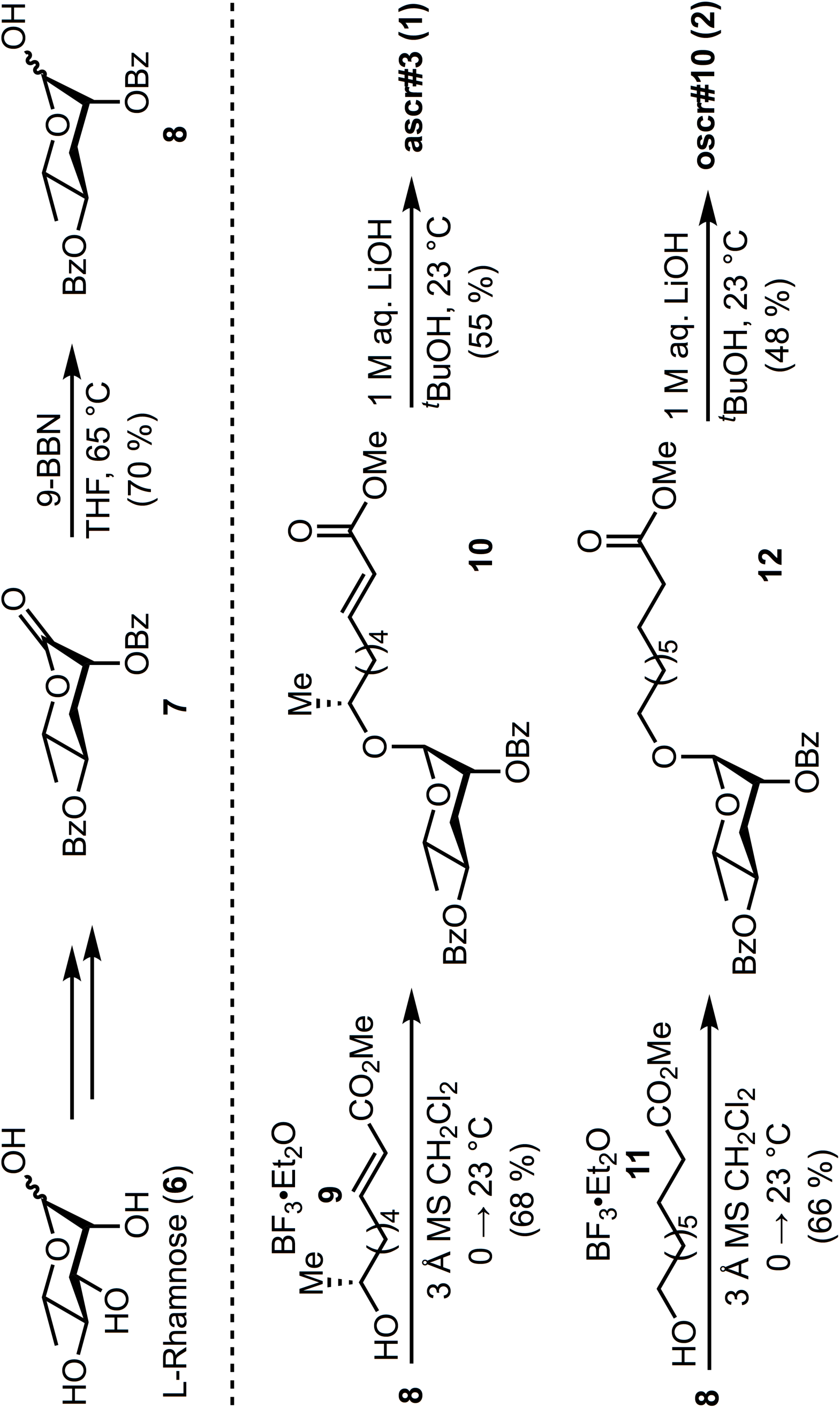
Synthesis of ascarosides. The synthetic pathway designed for protected ascarylose **8**, ascr#3 (**1**), oscr#10 (**2**) are illustrated.

## 3. Results

### 3.1. *Na*-ASP-2 binds cholesterol and palmitic acid *in vitro*

The *in vitro* cholesterol-binding activity of *Na*-ASP-2 was examined using increasing concentrations of radiolabeled [^3^H]-cholesterol and a constant concentration of purified protein. *Na*-ASP-2 displayed saturable binding of cholesterol with an apparent dissociation constant K_d_ of 2.1 µM (Figure 3A). *Na*-ASP-2 has similar cholesterol binding affinity as reported for other SCP/TAPS family members from yeast, *Saccharomyces cerevisiae* (Pry1, 1.9 µM), *Brugia malayi* (*Bm*-VAL-1, 0.9 µM), *Heligmosomoides polygyrus* (Hp-VAL-4, 1.53 µM) and *Schistosoma mansoni* (*Sm*-VAL-4, 2.4 µM) (Asojo et al., 2018; Darwiche et al., 2016; Darwiche et al., 2018; Kelleher et al., 2014). Furthermore, addition of equimolar or excess concentration of unlabeled cholesterol reduced binding of the radioligand, indicating that binding is specific as shown in Figure 3A,B.

**Figure 3.**
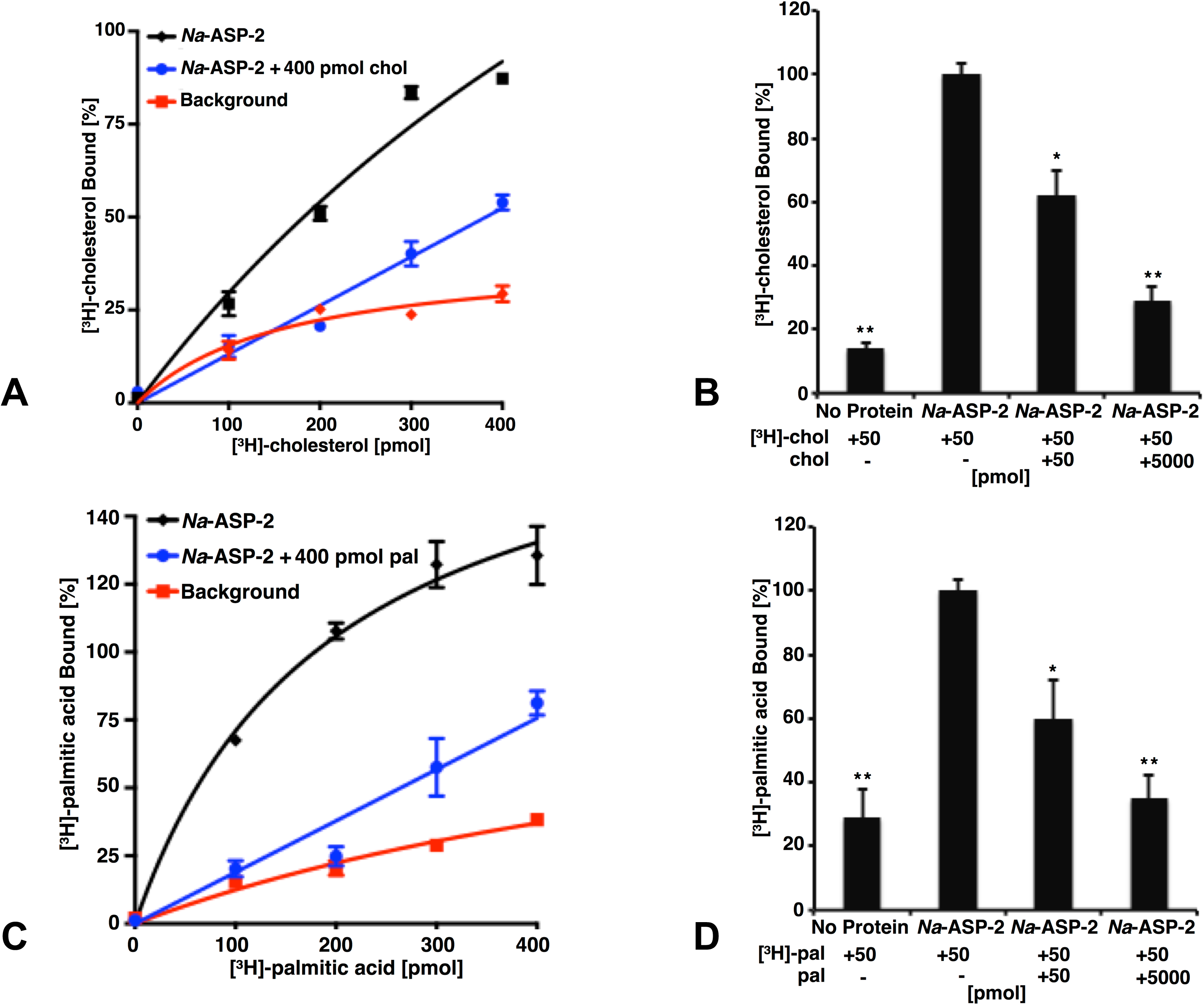
*Na*-ASP-2 binds both cholesterol and free palmitic acid. (A) Ligand binding of [^3^H]-cholesterol to *Na*-ASP-2. Data represent mean ± SD of 3 independent experiments. (B) Competitive binding of unlabeled cholesterol (50 or 5000 pmol) to *Na*-ASP-2. Each data point is the average of duplicate assays and represents the amount of [^3^H]-cholesterol bound relative to a control containing no unlabeled cholesterol. (C) Ligand binding of [^3^H]-palmitic acid to *Na*-ASP-2. (D) Competitive binding of unlabeled palmitic acid (50 or 5000 pmol) to *Na*-ASP-2. Each data point is the average of duplicate assays and represents the amount of [^3^H]-palmitic acid bound relative to a control containing no unlabeled palmitic acid. Data represent mean ± SD of 3 independent experiments. Asterisks denote statistical significance relative to the control containing only the radiolabeled ligand and either purified *Na*-ASP-2 or Pry1. (**, p < 0.001; *, p < 0.01).

Tablysin-15, a horsefly SCP/TAPS protein was shown to bind fatty acids with a hydrophobic pocket formed between two central helices (Ma et al., 2011). This hydrophobic pocket is observed in other SCP/TAPS proteins and we previously confirmed the ability of these proteins to bind palmitic acid *in vitro* (Asojo et al., 2018; Darwiche et al., 2016; Darwiche et al., 2018; Kelleher et al., 2014). To examine whether *Na*-ASP-2 can bind palmitic acid, we carried out direct binding studies using [^3^H]-palmitic acid as radiolabeled ligand, as shown in Figure 3C. *Na*-ASP-2 showed a saturable binding for palmitic acid with an apparent K_d_ of 95 µM, which is of the same magnitude as previously measured for the SCP/TAPS family members from yeast (Pry1, *K*_*d*_ =112 µM), *Brugia malayi* (*Bm*-VAL-1, *K*_*d*_ = 83 µM), and comparable to tablysin-15 (*K*_*d*_ = 94 µM) (Asojo et al., 2018; Darwiche et al., 2016; Darwiche et al., 2018; Kelleher et al., 2014). For competition binding assays, binding of *Na*-ASP-2 to palmitic acid was reduced in the presence of unlabeled palmitic acid, indicating that binding is specific (Figure 3C, D). Taken together our results indicate that *Na*-ASP-2 binds cholesterol and palmitic acid *in vitro*.

### 3.2. Fatty acids and ascaroside binding is selective for the palmitate-binding cavity

Having confirmed the ability of *Na-*ASP-2 to bind cholesterol, we carried out competitive binding studies of ascarosides and their fatty acid moieties against radiolabeled cholesterol. At a concentration of 50 pmol, the typical concentration for our cholesterol binding assay, neither ascarosides (ascr#3 (**1**) and oscr#10 (**2**)) nor fatty acids (**3-5**) competed with the radiolabelled [^3^H]-cholesterol (50 pmol) for binding to *Na*-ASP-2 (Figure 4A). We also tested if the ascarosides or their fatty acid moieties bind to the fatty acid binding cavity. Our studies showed that the binding of [^3^H]-palmitic acid by *Na*-ASP-2 was competed by the ascaroside, ascr#3 (**1**) and by all the fatty acid moieties **3**-**5** tested with the same order of magnitude, but not by the ascaroside, oscr#10 (**2**) (Figure 4B). We tested the ability of Pry1, a SCP/TAPS protein from *S. cerevisiae*, an organism that does not contain ascarosides, to bind to the same ligands. Our analysis revealed that while the fatty acids (**3-5**) competed for palmitic acid binding to Pry1, neither ascr#3 (**1**) nor oscr#10 (**2**) bound to Pry1. Furthermore, addition of excess ligands (fatty acids (**3-5**)) competed with radioligand binding while binding of [^3^H]-palmitic acid to Pry1 could not be competed for by the addition of excess unlabeled ascr#3 (**1**) or oscr#10 (**2**) (Figure 4C). We independently validated the binding of ligands to Pry1 and *Na*-ASP-2 by microscale thermophoresis and determined binding constants (Figure 5). The results of these analyses confirmed that Pry1 does not bind ascr#3 or oscr#10, but it binds palmitic acid and the fatty acid moieties present in ascarosides. *Na*-ASP-2, on the other hand, bound ascr#3 (**1**) with a K_d_ of 142 µM but did not bind oscr#10 (**2**), which is consistent with the results obtained by the ligand competition assay and indicates that *Na*-ASP-2 binds ascr#3 (**1**) through its fatty-acid binding pocket.

**Figure 4.**
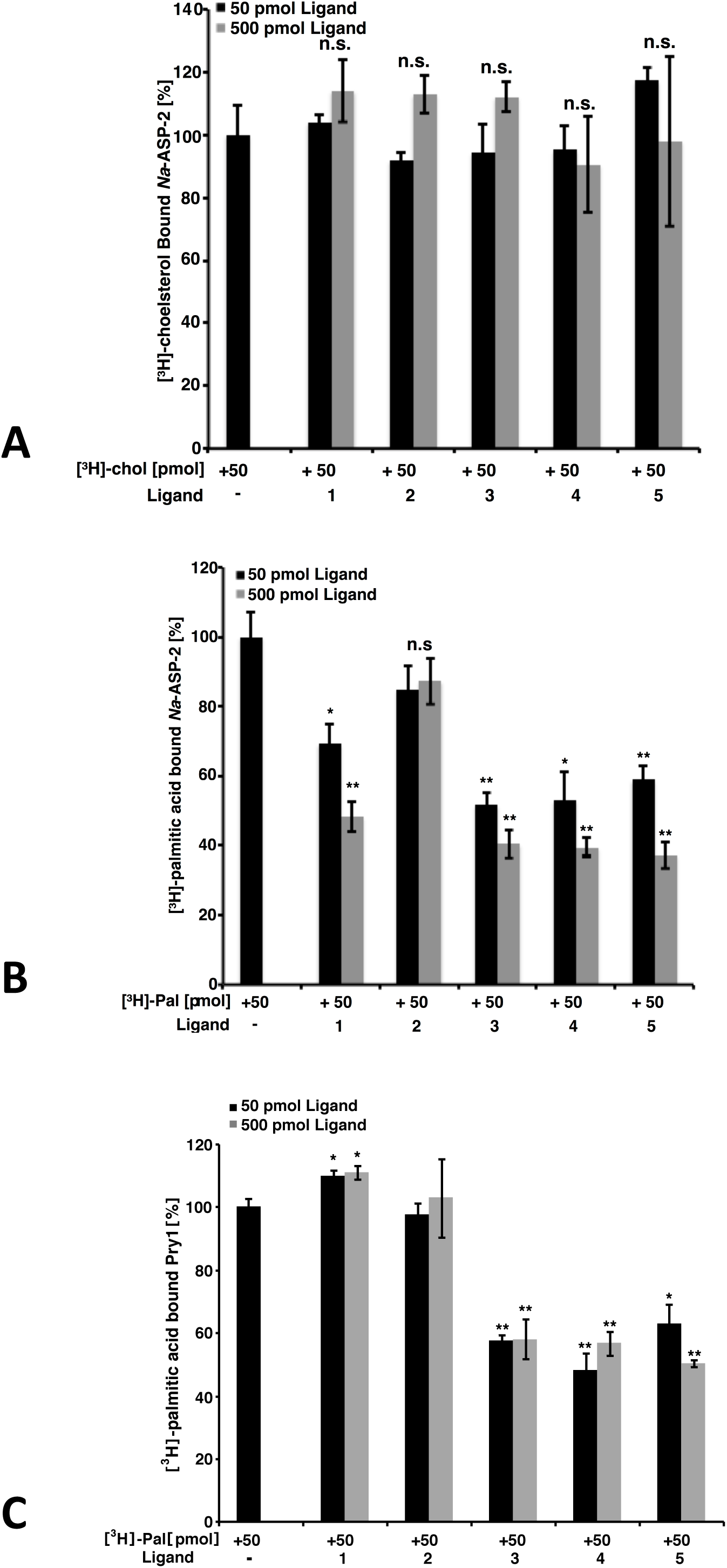
Binding of ascarosides to *Na*-ASP-2 and Pry1. (A) Free fatty acids and ascarosides fail to compete with [^3^H]-cholesterol for binding to *Na*-ASP-2. (B) Free fatty acids and ascarosides compete with [^3^H]-palmitic acid for binding to *Na*-ASP-2. (C) Free fatty acids but not ascarosides compete [^3^H]-palmitic acid for binding to Pry1. Competitive binding was tested with either 50 or 500 pmol of the unlabeled ligands and 50 pmol of [^3^H]-palmitic acid for binding to 100 pmol purified *Na*-ASP-2 or Pry1. The ascarosides tested are (**1)** (ascr#3) and (**2)** (oscr#10) while the fatty acids are **3** ((R)-8-hydroxynonanoic acid), **4** ((R,E)-8-hydroxynon-2-enoic acid), and **5** (9-hydroxynonanoic acid). Data represent mean ± SD of 3 independent experiments. Asterisks denote statistical significance relative to the control containing only the radiolabeled ligand and either purified *Na*-ASP-2 or Pry1. (**, p < 0.001; *, p < 0.01). n.s.; not significant.

**Figure 5.**
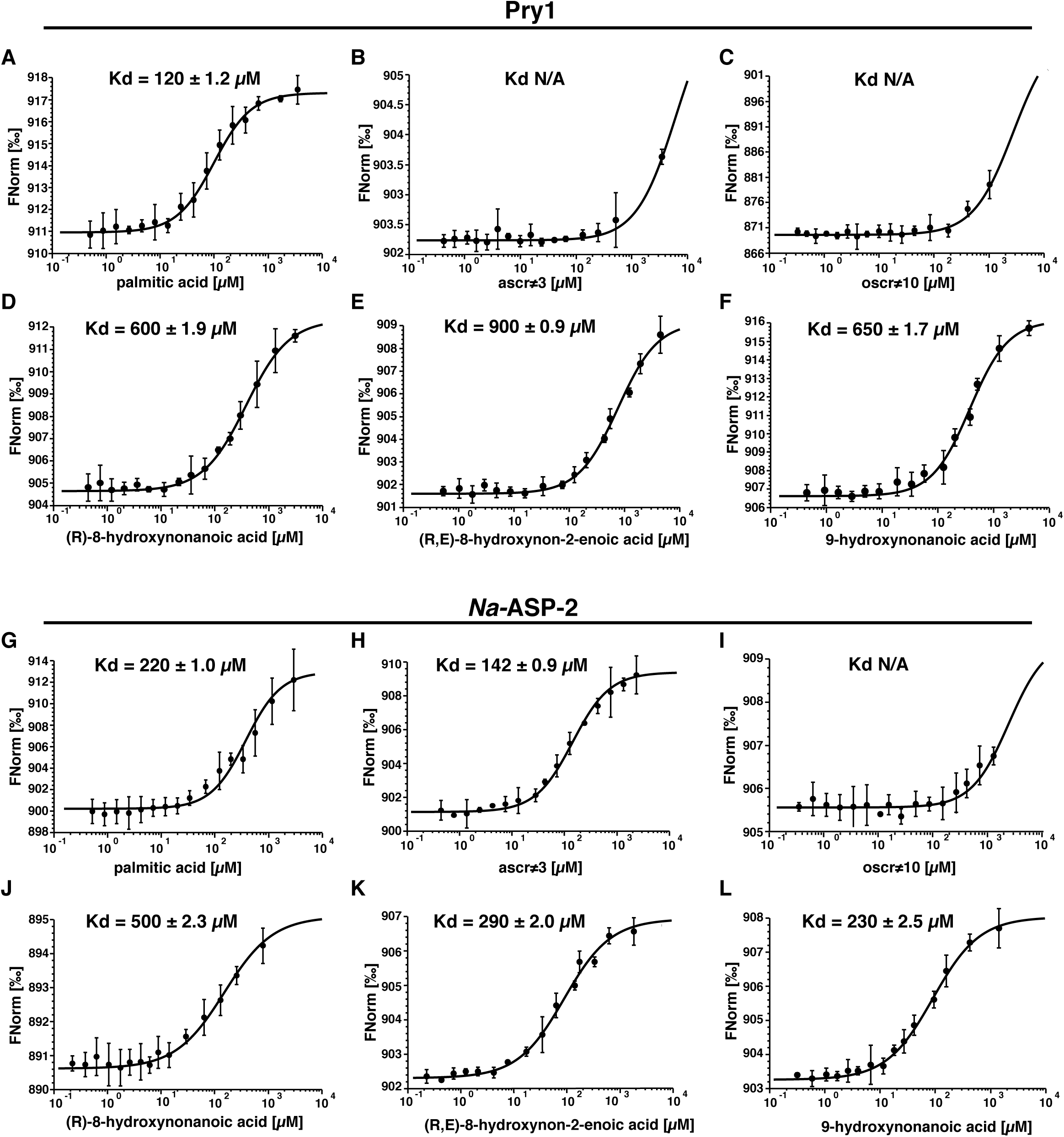
*Na*-ASP-2 selectively binds ascr#3 but not oscr#10. Binding of ascarosides and their fatty acid moieties by Pry1 and *Na*-ASP-2 as measured by microscale thermophoresis. (A, G) Palmitic acid; (B, H) ascr#3; (C, I) oscr#10; (D, J) (R)-8-hydroxynonanoic acid; (E, K) (R,E)-8-hydroxynon-2-enoic acid; (F, L) 9-hydroxynonanoic acid. Pry1 binds palmitic acid and free hydroxylated nanonoic acids with similar affinities but binds neither the ascarosides ascr#3 and oscr#10. Na-ASP-2 binds palmitic acid, ascr#3 and free hydroxylated nanonoic acids with similar affinities but not oscr#10. The Kd values are indicated in each figure with NA (not applicable) where there is no binding.

## 4. Discussion

We present here efficient methods to synthesize the ascarocides and their fatty acid moieties. We also present data revealing that the fatty acid moieties of ascarosides compete for binding to the palmitate-binding cavities of both Pry1 and *Na*-ASP-2 but as expected do not bind to the sterol binding cavity. The micromolar binding affinity of ascr#3 and free fatty acids are comparable to that observed for palmitic acid to the palmitate-binding cavity of other CAP proteins. While it is unclear if ascr#3 binding is physiologically relevant, the finding that ascr#3 (**1**) binds *Na*-ASP-2 is still interesting considering that a high relative abundance of ascr#3 (1) was detected in E-S products from both the infective juvenile and adult stages of *Nippostrongylus brasilensis* by HPLC-MS(Choe et al., 2012). It is plausible that ascr#3 (**1**) is present in human hookworms since there appears to be a conservation of ascaroside production in families of nematodes (Choe et al., 2012). A blast search of the *Na*-ASP-2 sequence against the *N. brasilensis* proteins reveals several SCP/TAPs proteins, which share over 45 % sequence similarity with *Na*-ASP-2. Even more remarkable, the residues and predicted structures of the helical regions notably residues corresponding to (alpha 1 and alpha 3) that form the fatty acid-binding cavity are conserved (Figure 6A). This structural similarity suggests that these proteins likely behave similarly to *Na*-ASP-2 as we observed previously for the orthologues from *B. malayi* and *H. polygyrus*. Additionally, we observed that the incorporation of the ascarylose sugar abrogated the ability of these fatty acids to bind to Pry1. A comparison of the helices bordering the palmitic acid binding cavities of Pry1 and *Na*-ASP-2 reveals that Pry1 has shorter helices than *Na*-ASP-2, which results in a smaller hydrophobic binding pocket in Pry1 compared to *Na*-ASP-2 (Figures 6A and B). This smaller size may explain the failure of Pry1 to accommodate ascaroside as opposed to free fatty acids. The inability of *Na*-ASP-2 to bind oscr#10 (**2**) cannot be explained by the size difference of the cavities and suggests a new hypothesis that we plan to test in future; that ascaroside binding may be specific for certain SCP/TAPS proteins, indicating a possible functional relationship between ascarosides and parasite SCP/TAPS proteins.

**Figure 6.**
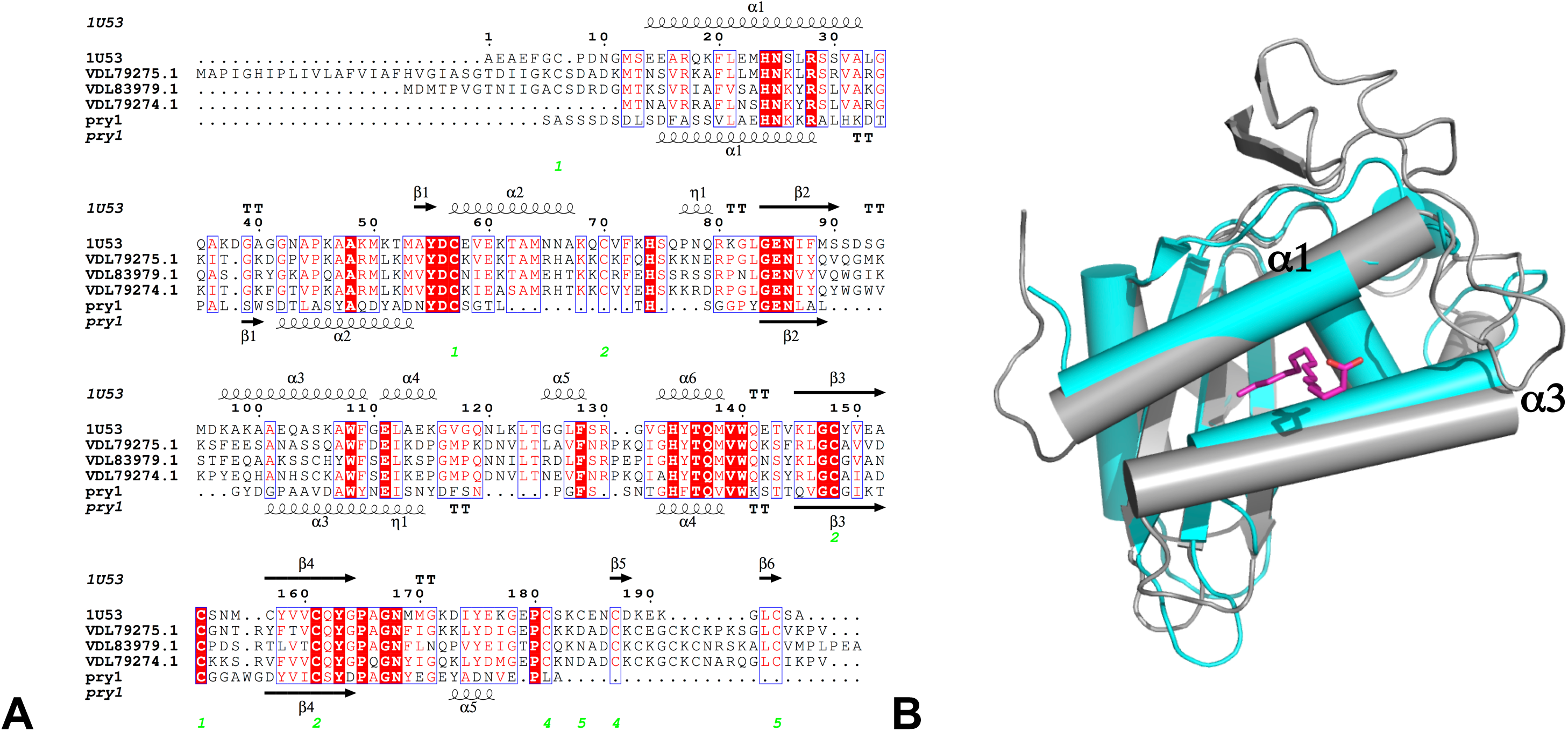
Comparison of fatty acid binding cavities of *Na*-ASP-2 and Pry1. (A) Structure based alignment of *Na*-ASP-2, Pry1 and three *N. brasilensis* SCP/TAPs proteins (genbank codes VDL79275.1; VDL83979.1; and VDL79274.1). The sequences are aligned with clustalWOmega and the secondary structural features are illustrated with the coordinates of HpVAL-4 and Pry1 using ESPript. (Gouet et al., 2003) The alpha helices (alpha 1 and alpha 3) that form the palmitate-binding cavity have similar lengths for *Na*-ASP-2 and the *N. brasilensis* proteins whereas Pry1 has shorter helices. The secondary structure elements shown are alpha helices (α), 3_10_-helices (η), beta strands (β), and beta turns (TT). Identical residues are shown in solid red, and conserved residues are in red. The locations of the cysteine residues involved in disulfide bonds are numbered in green. (B) Both of the helices (α1 and α 3) forming the palmitic acid binding cavity of Pry1 (cyan) are shorter than those from *Na*-ASP-2 (gray). Also shown in magenta is the stick structure of palmitate superposed from the structure of the complex of tablysin-15 with palmitate.

## 5. Conclusions

In summary, our results reveal that the fatty acid moieties of the ascarosides, ascr#3 (**1**) and oscr#10 (**2**), bind specifically to the fatty acid binding cavity of both *Na*-ASP-2 and Pry1, with the latter protein from *Saccharomyces cerevisiae* SCP/TAPS serving as a control. Additionally, ascr#3 (**1**), an ascaroside that is present in mammalian hookworm E-S products binds competitively to the fatty acid binding cavity of *Na*-ASP-2, whereas oscr#10 (**2**) which is not found in hookworm E-S products did not. Interestingly, neither ascaroside bound to Pry1. Studies to identify how ascarosides precisely interact with parasite CAP proteins are currently underway. More studies need to be conducted to determine the physiological relevance of the fatty acid-binding cavity of *Na*-ASP-2.

## Supporting information

see SI for synthetic details

## Acknowledgments

RS thanks the Swiss National Science Foundation for support (grant 31003A_173003). This study was supported in part by funds from the Collaborative Faculty Research Investment Program of Baylor University, Baylor Scott & White Health and Baylor College of Medicine to OAA and DR.

## Author Contributions Statement

Designed the studies: OAA, OEA, RS, RD, KDH, DR

Conducted experiments: OEA, RD, NJT

All authors contributed expertise and to the final manuscript.

## Notes

### Competing Interest Statement

The authors have declared no competing interest.

