## Supplementary material for "Necator americanus Ancylostoma secreted protein-2 (Na-ASP-2) selectively binds an ascaroside (ascr#3)": see SI for synthetic details

**Supplementary data for *Necator americanus Ancylostoma* secreted protein-2 (*Na*-ASP-2) selectively binds an ascaroside (ascr#3)**

Ola El Atab[1+], Rabih Darwiche[1,2+], Nathanyal J. Truax[3], Roger Schneiter[1], Kenneth G. Hull[3], Daniel Romo[3] and Oluwatoyin A. Asojo[4,5*]

[1]Division of Biochemistry, Department of Biology, University of Fribourg, Chemin du Musée 10, CH 1700 Fribourg, Switzerland
[2]Department of Biological Chemistry and Molecular Pharmacology, Harvard Medical School, Boston, MA 02115
[3]Department of Chemistry and Biochemistry & The CPRIT Synthesis and Drug-Lead Discovery Laboratory, Baylor University, One Bear Place, Waco, Texas 76798-7348, United States
[4]Department of Chemistry and Biochemistry, 200 William Harvey Way, Hampton University, Hampton VA, 23668
[5]National School of Tropical Medicine, Baylor College of Medicine, Houston Texas, 77030

### Part 1: Synthesis
### General

$^1$H NMR and $^{13}$C NMR were recorded at 25 °C using either a 600 equipped with Prodigy Cold Probe NMR ($^1$H NMR at 600 MHz, $^{13}$C NMR at 150 MHz), 500 ($^1$H NMR at 500 MHz, $^{13}$C NMR at 125 MHz), or 400 ($^1$H NMR at 400 MHz, $^{13}$C NMR at 100 MHz). Chemical shifts are reported in ppm using the solvent resonance as the internal standard ($^1$H NMR $CDCl_3$: δ 7.26 ppm, $^{13}$C NMR $CDCl_3$: δ 77.16 ppm, $^1$H NMR $CD_3OD$: δ 3.31 ppm $^{13}$C NMR $CD_3OD$: δ 49.0 ppm) Data is reported as follows: chemical shift, multiplicity (s = singlet, d = doublet, t = triplet, q = quartet, bs = broad singlet, m = multiplet (or any combination of these), app = apparent when multiplicity arises from coincidental equivalence of coupling constants, or there is obviously higher-order coupling than can be resolved within a given resonance *e.g.* app t = apparent triplet), coupling constants (Hz) and integration. Infrared spectra (IR) were obtained using both ATR and thin film (NaCl plates) sampling techniques were used) and recorded in wavenumbers ($cm^{-1}$). Bands are characterized as broad (br), strong (s), medium (m), and weak (w). High Resolution Mass Spectrometry (HRMS) analysis was obtained using Thermo Orbitrap Discovery utilizing Electrospray Ionization (ESI) and are reported as m/z (relative intensity). Optical rotations were recorded at 589 nm employing a 25 mm cell. Specific rotations $[\alpha]_D^{25}$, are reported in degree mL/(g•dm) at the specific temperature.

All non-aqueous reactions were performed under a nitrogen atmosphere in oven-dried (125 °C) or flame-dried glassware unless otherwise indicated. Reaction solvents used were pre-dried by passing through activated molecular sieves or alumina in a JC Meyer Solvent Drying System. Both diisopropylethylamine (DIPEA) and triethylamine ($Et_3N$) were distilled over $CaH_2$ prior to use. All work-up and purifications were completed using reagent grade solvents and no precautions were taken to exclude air. Thin Layer Chromatography (TLC) was performed using glass-backed silica gel F254 (Silicycle, 250 µm thickness). TLCs were visualized under UV irradiation (254 nm) or by the use of Hanessian's or $KMnO_4$ staining solutions as specified for each reaction. Standard Flash column chromatography was completed using Silicycle ultrapure SiliaFlash silica gel, 40-63 µm, 60 Å pore size. As specified, in some cases medium pressure liquid chromatography

(MPLC) was performed using a Teledyne Isco CombiFlash Rf automated flash chromatography system.

**Experimental Procedures**

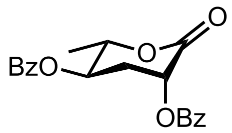

7

**(2*S*,3*R*,5*R*)-2-methyl-6-oxotetrahydro-2*H*-pyran-3,5-diyl dibenzoate (7):** Synthesized in a 5 step sequence as previously reported by Paik and co-workers.[1] Characterization data matched that previously reported.

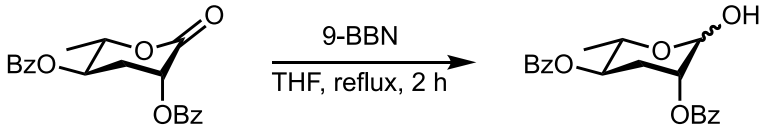


7 8

**(2*S*,3*R*,5*R*)-2-methyl-6-oxotetrahydro-2*H*-pyran-3,5-diyl dibenzoate (8):** The synthetic route to this intermediate is a modified procedure from that previously reported by Paik and co-workers.[1] An oven dried 25 mL RB flask was charged with lactone **7** (200 mg, 0.564 mmol, 1.0 equiv.). The RB was then equipped with a reflux condenser and the entire apparatus was backfilled with $N_2$ (3x) and the solid was taken up in THF (5.6 mL). To the resulting solution, stirring at ambient temperature (23 °C), was added dropwise, a 0.5 M solution of 9-BBN in THF (3.4 mL, 1.69 mmol, 3.0 equiv.). The solution was then heated to reflux (65 °C) and was allowed to stir. After competition as judged by TLC (2 h), the reaction mixture was cooled to 0 °C and then treated with 35 % $H_2O_2$ (4 mL). The bulk of the THF was removed in vacuo and the remaining aqueous layer was extracted with $CH_2Cl_2$ (2 x 10 mL). The combined organic extracts were dried over $Na_2SO_4$, filtered, and the solvent was removed in vacuo to yield a clear colorless sticky oil. The oil was purified by MPLC (4 g silica, 0 → 15 % EtOAc in $CH_2Cl_2$) to yield protected ascarylose **8** (141 mg, 0.394 mmol, 70 %). Spectral data for this compound was consistent with that previously reported.[1] TLC (10:2 toluene:EtOAc v/v, UV) $R_f$ = 0.47; NMR data is provided here since data was taken at a higher field strength than previously reported. $^1$H NMR (400 MHz, $CDCl_3$): δ 1.59 (d, *J* = 6.5 Hz, 3H), 1.84-1.91 (m, 1H), 2.39-2.42 (m, 1H), 2.58 (ddd, *J* = 14.3, 7.5, 3.6 Hz, 1H), 2.71 (ddd, *J* = 14.1, 12.1, 6.1 Hz, 1H), 4.82 (pent., *J* = 6.4 Hz, 1H), 5.28 (dt, *J* = 6.0, 3.6 Hz, 1H), 5.90, (dd, *J* = 12.1, 7.5 Hz, 1H), 7.44-7.51 (m, 4H), 7.58-7.65 (m, 2H), 8.05-8.11 (m, 4H).

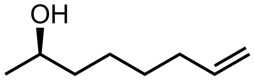

13

**(*R*)-oct-7-en-2-ol (13):** Synthesized as previously reported by Paik and co-workers.[1] Characterization data matched that previously reported.

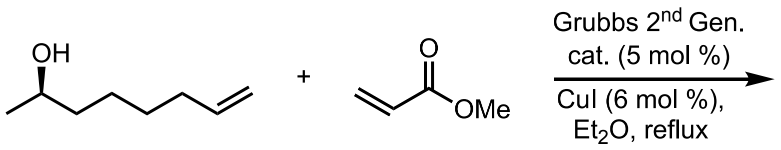


13 9

**Methyl (8*R*,2*E*)-8-hydroxynon-2-enoate (9):** Adapting a previously reported method,[2] an oven dried RB flask was charged with Grubbs 2nd generation catalyst (166 mg, 0.190 mmol, 0.05 equiv.) and CuI (50.0 mg, 0.263 mmol, 0.067 equiv.). The RB flask was then equipped with a reflux condenser and then backfilled with Argon (3x) and the solid was taken up in $Et_2O$ (3.9 mL). To the stirred suspension was added methyl acrylate (1.8 mL, 20 mmol, 5.0 equiv.). The reaction mixture was allowed to stir for 10 min prior to the addition of chiral alcohol (**S1**) (500 mg, 3.90 mmol, 1.0 equiv.). The vial was then sealed and the mixture was then heated to 40 °C and stirred until complete conversion was noted by TLC (~ 4h). The solvent was then removed in vacuo to give a black crude material that was then purified by MPLC (12 g silica, 0 → 50 % EtOAc in hexanes) to yield a dark oil which was purified again by MPLC (12 g silica, 0 → 30 % EtOAc in $CH_2Cl_2$) to yield the ester **9** as a clear, light brown, non-viscous oil (531 mg, 2.85 mmol, 71 % yield). TLC (1:2 EtOAc:hexanes v/v, $KMnO_4$ Stain) $R_f$ = 0.30; $[\alpha]_D^{25.3}$ – 9.05 (*c* = 1.41, MeOH); $^1H$ NMR (400 MHz, $CDCl_3$): δ 1.19 (d, *J* = 6.2 Hz, 3H), 1.53-1.29 (m, 7H), 2.22 (app dq, *J* = 7.1, 1.4 Hz, 2H), 3.72 (s, 3H), 3.84-3.75 (m, 1H), 5.82 (dt, *J* = 15.6, 1.6 Hz, 1H), 6.96 (dt, *J* = 15.6, 7.0 Hz, 1H) ppm; $^{13}C$ NMR (100 MHz, $CDCl_3$): δ 23.7, 25.4, 28.2, 32.3, 39.2, 51.6, 68.1, 121.2, 149.6, 167.3; IR (thin film): 3418 (br, m), 2933 (m), 2859 (m), 1728 (m-s), 1658 (m), 1436 (m) $cm^{-1}$; HRMS (ESI+) *m/z* calcd for $C_{10}H_{18}O_3$ $[M + Na]^+$: 209.1148; found: 209.1149.

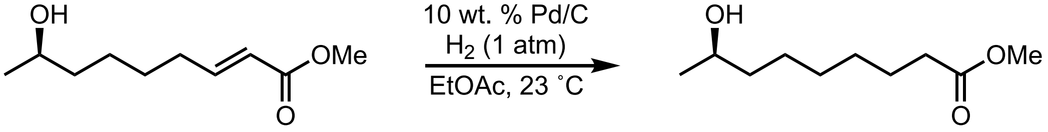


9 14

**Methyl (*R*)-8-hydroxynonanoate (14):** To a RB flask containing 10 wt % Pd/C (wet, 82 mg, 0.08 mmol, 0.10 equiv.) under $N_2$, was added a solution of enoate **9** (145 mg, 0.779 mmol, 1.0 equiv.) in EtOAc (3.9 mL). The head space was then backfilled with $H_2$ (3x) using a $H_2$ filled balloon. The reaction was allowed to stir under an atmosphere of $H_2$ at ambient temperature (23 °C) until the starting material was fully consumed (as judged by TLC, ~3.5 h). The reaction mixture was then filtered through celite and the solvent was removed in vacuo to yield the title compound as a light yellow, non-viscous oil (138 mg, 0.733 mmol, 94 %) which was of sufficient purity to be used in the next step without further purification. Spectral data for this compound was consistent with that previously reported.[3] TLC (2:1 EtOAc:hexane v/v, $KMnO_4$) $R_f$ = 0.48; $^1H$ NMR (500 MHz, $CDCl_3$): δ 1.18 (d, *J* = 6.19 Hz, 3H), 1.27-1.48 (m, 9H), 1.56-1.67 (m, 2H), 2.30 (t, *J* = 7.5 Hz, 2H), 3.66 (s, 3H), 3.74-3.83 (m, 1H); $^{13}C$ NMR (125 MHz, $CDCl_3$): δ 23.7, 25.0, 25.7, 29.2, 29.4, 34.2, 39.4, 51.6, 68.2, 174.4.

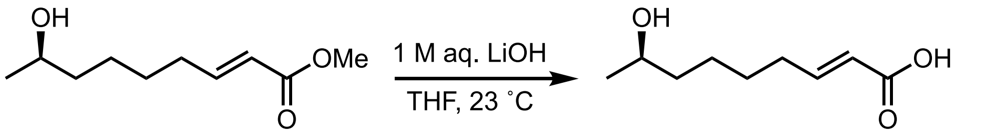


9 3

**(8*R*,2*E*)-8-Hydroxynon-2-enoic acid (3):** To a 23 °C stirred solution of chiral enoate **9** (60.0 mg,

0.322 mmol, 1.0 equiv.) in THF (5 mL), was added a 1M aqueous LiOH solution (5 mL). The reaction was allowed to stir at 23 °C until full consumption of starting material was observed by TLC (~4.5 h). The pH was then adjusted to ~4 using 1M HCl. The biphasic mixture was then separated and the aqueous layer was extracted with EtOAc (2x 5 mL). The combined organic layers were dried over $MgSO_4$ and the solvent was removed in vacuo. The crude material was then purified by flash chromatography (0 → 10 % MeOH in $CH_2Cl_2$) to yield a light-yellow oil. This material was then purified again by flash chromatography (0 → 3 % MeOH in $CH_2Cl_2$) to yield the title compound as a colorless solid film (30.0 mg, 0.174 mmol, 54 %). TLC (1:1 EtOAc:hexane v/v, $KMnO_4$) $R_f$ = 0.09; $[\alpha]_D^{24.4}$ – 8.49 (*c* = 0.47, MeOH); $^1H$ NMR (600 MHz, $CDCl_3$): δ 1.19 (d, *J* = 6.1 Hz, 3H), 1.31-1.55 (m, 6H), 2.24 (app q, *J* = 7.2 Hz, 2H), 3.77-3.84 (m, 1H), 5.42-6.09 (m, RCOOH, ROH, alkene CH, 3H), 7.05 (dt, *J* = 14.6, 6.9 Hz, 1H); $^{13}C$ NMR (150 MHz, $CDCl_3$): δ 23.5, 25.4, 28.0, 32.4, 39.1, 68.2, 121.1, 151.8, 171.6; IR (ATR) 3371 (br, w), 2935 (m), 2860 (w), 1684 (s), 1652 (m), 1636 (m), 1283 (m) $cm^{-1}$; HRMS (ESI+) *m/z* calcd for $C_9H_{16}O_3$ $[M + Na]^+$: 195.0992; found: 195.0993.

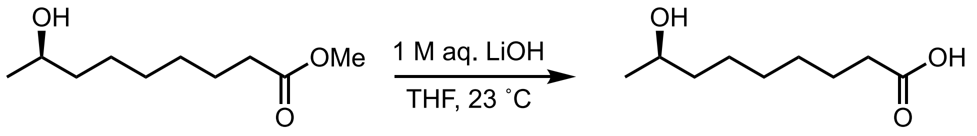


14 4

**(*R*)-8-Hydroxynonanoic acid (4):** To a 23 °C stirred solution of ester **S2** (50.0 mg, 0.266 mmol, 1.0 equiv.) in THF (5 mL), was added a 1M aqueous LiOH solution (5 mL). The reaction was allowed to stir at 23 °C until full consumption of starting material was observed by TLC. The pH was then adjusted to ~3 using 1M HCl. The bulk of the THF was removed in vacuo and the aqueous layer was extracted with $Et_2O$ (2x 10 mL). The combined organic layers were washed with brine and dried over $Na_2SO_4$, filtered, and concentrated in vacuo to yield a colorless oil that was then purified by flash chromatography (0 → 70 % EtOAc in hexanes) to yield the title compound as a waxy solid (23.0 mg, 0.132 mmol, 50 %). TLC (EtOAc, Hanessian's stain) $R_f$ = 0.40; $[\alpha]_D^{24.4}$ – 6.39 (*c* = 0.31, MeOH); $^1H$ NMR (400 MHz, $CDCl_3$): δ 1.17 (d, *J* = 6.1 Hz, 3H) 1.25-1.50 (m, 8H), 1.57-1.67 (m, 2H), 2.32 (t, *J* = 7.5 Hz, 2H), 3.75-3.83 (m, 1H), 6.08 (bs, 1H, ROH); $^{13}C$ NMR (100 MHz, $CDCl_3$): δ 23.4, 24.7, 25.6, 29.1, 29.3, 34.1, 39.2, 68.3, 179.4; IR (ATR) 3347 (br, w), 2927 (m), 2856 (m-w), 1703 (s) $cm^{-1}$; HRMS (ESI+) *m/z* calcd for $C_9H_{18}O_3$ $[M + Na]^+$: 197.1148; found: 197.1150.

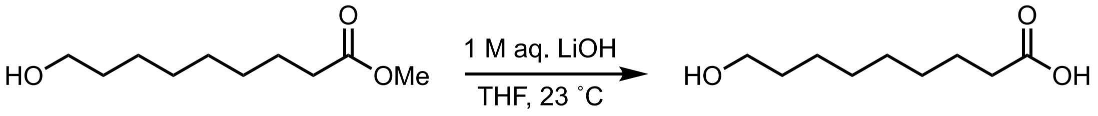


5

**9-Hydroxynonanoic acid (5):** To a 23 °C stirred solution of commercially available 9-hydroxypelargonic acid methyl ester **11** (65.0 mg, 0.345 mmol, 1.0 equiv.) in THF (5 mL), was added a 1M aqueous LiOH solution (5 mL). The reaction was allowed to stir at 23 °C until full consumption of starting material was observed by TLC. The pH was then adjusted to ~3 using 1M HCl. The bulk of the THF was removed in vacuo and the aqueous layer was extracted with $Et_2O$ (2x 10 mL). The combined organic layers were washed with brine, dried over $Na_2SO_4$, filtered, and concentrated in vacuo to yield a colorless solid that was then purified by flash chromatography (0 → 50 % EtOAc in hexanes) to yield the title compound as a colorless oily solid (38.0 mg, 0.132 mmol, 50 %). TLC (EtOAc, Hanessian's stain) $R_f$ = 0.33; $^1H$ NMR (500 MHz, $CDCl_3$): δ 1.27-1.38 (m, 8H), 1.51-1.67 (m, 4H), 2.34 (t, *J* = 7.5 Hz, 2H), 3.64, (t, *J* = 6.6 Hz, 2H),

6.10 (bs, 2H, ROH and RCOOH); $^{13}$C NMR (100 MHz, $CDCl_3$): δ 24.8, 25.7, 29.1, 29.3 (2), 32.7, 34.2, 63.1, 179.6; IR (ATR) 3400 (br, w) 2930 (m), 2852 (m), 1688 (m-s) 1194 (m) $cm^{-1}$; HRMS (ESI+) *m/z* calcd for $C_9H_{18}O_3$ $[M + Na]^+$: 197.1148; found: 197.1150.

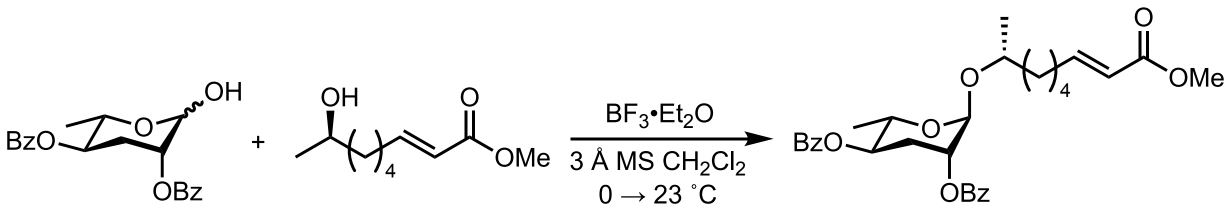


8 9 10

**(2*R*,3*R*,5*R*,6*S*)-2-(((*R*,*E*)-9-methoxy-9-oxonon-7-en-2-yl)oxy)-6-methyltetrahydro-2*H*-pyran-3,5-diyl dibenzoate (10):** A flame dried vial was charged with protected ascarylose **8** (100 mg, 0.281 mmol, 1.0 equiv.) and powdered 4 Å molecular sieves (13 mg). The vial was backfilled with $N_2$ (3x) and the mixture was taken up in $CH_2Cl_2$ (2.0 mL). The solution was then cooled to 0 °C and chiral alcohol **9** (68 mg, 0.365 mmol, 1.3 equiv) was added followed by $BF_3$•$Et_2O$ (69 µL, 0.561 mmol, 2.0 equiv.). The 0 °C ice bath was then removed and the reaction was allowed to stir and warm to ambient temperature (23 °C). After full consumption of protected ascarylose as determined by TLC (~18 h), $Et_3N$ (0.5 mL) was added and the suspension was filtered through celite. The solvent was then removed in vacuo to yield a light-yellow oil that was purified by MPLC (12 g silica, 0 → 50 % EtOAc in hexanes) to yield the title compound as a clear, colorless, viscous oil (100 mg, 0.191 mmol, 68 %). Spectral data for this compound was consistent with that previously reported[4] TLC (1:2 EtOAc:hexane v/v) $R_f$ = 0.51; $^1$H NMR (400 MHz, $CDCl_3$): δ 1.19 (d, *J* = 6.07 Hz, 3H), 1.28 (d, *J* = 6.25 Hz, 3H), 1.38-1.68 (m, 6H), 2.15-2.30 (m, 3H), 2.42 (app dt, *J* = 13.5, 4.0 Hz, 1H), 3.70 (s, 3H), 3.80-3.89 (m, 1H), 4.09 (m, 1H), 4.95 (s, 1H), 5.11-5.22 (m, 2H), 5.85 (dt, *J* = 15.7, 1.6 Hz, 1H), 7.00 (dt, *J* = 15.7, 7.0 Hz, 1H), 7.42-7.51 (m, 4H), 7.54-7.62 (m, 2H), 8.01-8.15 (m, 4H); $^{13}$C NMR (100 MHz, $CDCl_3$): δ 18.0, 19.3, 25.4, 28.1, 29.9, 32.3, 37.0, 51.6, 67.2, 70.8, 71.4, 72.7, 94.0, 121.2, 128.6, 129.8, 130.0, 130.1, 133.3, 133.4, 149.5, 165.8, 165.9, 167.3.

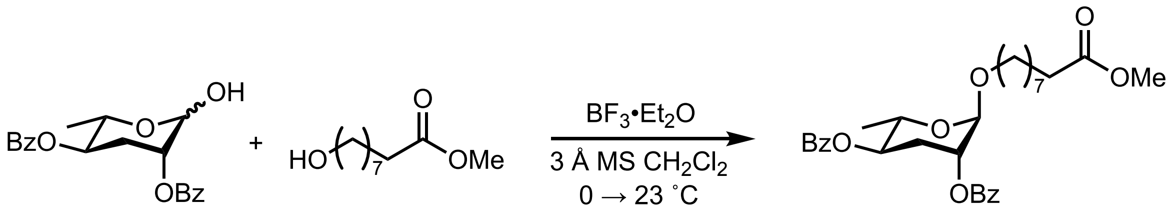


8 12

**9-(((2*R*,3*R*,5*R*,6*S*)-3,5-bis(benzoyloxy)-6-methyltetrahydro-2*H*-pyran-2-yl)oxy)nonanoic acid (12):** A flame dried vial was charged with protected ascarylose **8** (195.0 mg, 0.267 mmol, 1.0 equiv.) and powdered 4 Å molecular sieves (13 mg). The vial was backfilled with $N_2$ (3x) and the mixture was taken up in $CH_2Cl_2$ (2.0 mL). The solution was then cooled to 0 °C and commercially available 9-hydroxypelargonic acid methyl ester **11** (65 mg, 0.347 mmol, 1.3 equiv) was added followed by $BF_3$•$Et_2O$ (66 µL, 0.533 mmol, 2.0 equiv.). The 0 °C ice bath was then removed and the reaction was allowed to stir and warm to ambient temperature (23 °C). After full conversion of protected ascarylose as determined by TLC (~18 h), $Et_3N$ (0.5 mL) was added and the suspension was filtered through celite. The solvent was then removed in vacuo to yield a light yellow oil that was purified by MPLC (12 g silica, 0 → 50 % EtOAc in hexanes) to yield the title compound as a clear, colorless, viscous oil (93 mg, 0.177 mmol, 66 %). Spectral data for this compound was consistent with that previously reported.[5] TLC (1:4 EtOAc:hexane v/v) $R_f$ = 0.48; $^1$H NMR (500 MHz, $CDCl_3$): δ 1.28-1.44 (m, 11H), 1.58-1.69 (m, 4H), 2.21 (ddd, *J* = 13.5, 11.4, 3.2 Hz, 1H), 2.31 (t, *J* = 7.6 Hz, 2H), 2.41 (app dt, *J* = 13.5, 3.2 Hz, 1H), 3.50 (dt, *J* = 9.7, 6.5 Hz, 1H), 3.66 (s, 3H), 3.75 (dt, *J* = 9.7, 6.7 Hz, 1H), 4.07 (app dq, *J* = 9.7, 6.3 Hz, 1H), 4.82 (s, 1H), 5.14-5.22 (m, 2H), 7.42-7.50 (m, 4H), 7.55-7.60 (m, 2H), 8.02-8.06 (m, 2H), 8.09-8.13 (m, 2H); $^{13}$C NMR (125 MHz, $CDCl_3$): δ 18.0, 25.1, 26.22, 29.23, 29.3, 29.4, 29.9, 34.2, 51.6, 66.8, 68.0, 70.7, 96.5, 128.6, 129.7, 129.96, 130.00, 130.1, 133.3, 133.4, 165.8, 165.9, 174.4.

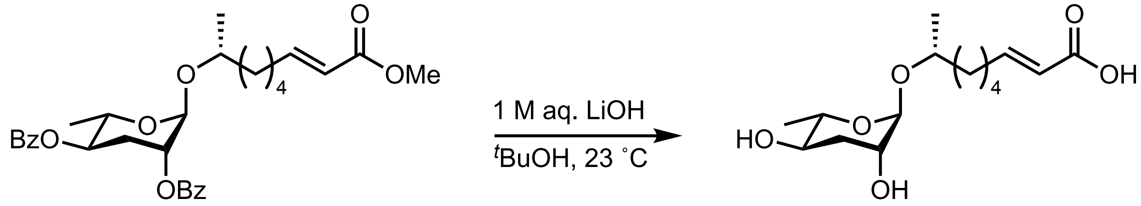

10 1

**(*R*,*E*)-5-(((2*R*,3*R*,5*R*,6*S*)-3,5-dihydroxy-6-methyltetrahydro-2*H*-pyran-2-yl)oxy)hex-2-enoic acid (ascr#3, 1):** To a stirred solution of protected ascr#3 **10** (95.0 mg, 0.181 mmol, 1.0 equiv), in $^t$BuOH (14 mL) at 23 °C, was added a 1M aqeuos LiOH solution (14 mL). The reaction was allowed to stir at 23 °C until full conversion of starting material was apparent by TLC (~18h). 1M HCl was then added until a pH of ~3 was achieved. The solution was then extracted with $^i$PrOH/$CH_2Cl_2$ (1:4 v/v) (3 x 20 mL). The aqueous layer was then saturated with NaCl, and extracted with $^i$PrOH/$CH_2Cl_2$ (1:4 v/v) (4 x 20 mL) and the combined organic layers were dried over $Na_2SO_4$, filtered, and the solvent was removed in vacuo to yield a colorless solid that was purified by flash chromatography (0 → 3 % MeOH in $CH_2Cl_2$) to yield ascr#3 (**1**) as a clear colorless film (30.0 mg, 0.099 mmol, 55 % yield). Spectral data for this compound was consistent with that previously reported.[4] TLC (1:9 MeOH:$CH_2Cl_2$ v/v, Hanessian's stain) $R_f$ = 0.27; $^1$H NMR (600 MHz, $CD_3OD$) δ 1.12 (d, *J* = 6.1 Hz, 3H), 1.21 (d, *J* = 6.2 Hz, 3H), 1.39-1.61 (m, 6H), 1.76 (ddd, *J* = 13.7, 11.5, 3.2 Hz, 1H), 1.95 (dt, *J* = 13.1, 4.0 Hz, 1H) 2.24 (aq, *J* = 6.7 Hz, 2H), 3.52 (ddd, *J* = 11.3, 9.4, 4.6, 1H), 3.59-3.65 (m, 1H), 3.71 (as, 1H), 3.75-3.82 (m, 1H), 4.64 (s, 1H), 5.81 (d, *J* = 15.5 Hz, 1H), 6.94 (dt, *J* = 14.9, 7.0 Hz, 1H); $^{13}$C NMR (150 MHz, $CD_3OD$): δ 18.1, 19.3, 26.4, 29.2, 33.1, 36.0, 38.1, 68.3, 70.0, 71.2, 72.4, 97.6, 122.9, 150.7, 170.4; IR (thin film) 3366 (br, m), 2933 (m), 1698 (m), 1653 (w-m), 1028 (s) cm$^{-1}$.

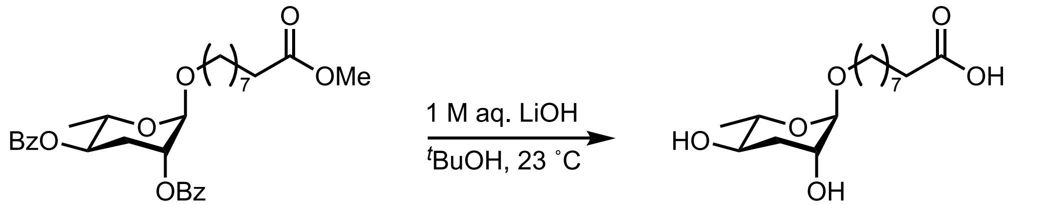

12 2

**9-(((2*R*,3*R*,5*R*,6*S*)-3,5-dihydroxy-6-methyltetrahydro-2*H*-pyran-2-yl)oxy)nonanoic acid (oscr#10, 2):** To a 23 °C stirred solution of protected oscr#10 **12** (98.0 mg, 0.177 mmol, 1.0 equiv.), in $^{t-}$BuOH (14 mL), was added a 1M aqueous LiOH solution (14 mL). The reaction was allowed to stir at 23 °C until full conversion of starting material was apparent by TLC (~18h). 1M HCl was then added until a pH of ~3 was achieved. The solution was then extracted with $^i$PrOH/$CH_2Cl_2$ (1:4 v/v) (3 x 20 mL). The aqueous layer was then saturated with NaCl, and extracted with $^i$PrOH/$CH_2Cl_2$ (1:4 v/v) (4 x 20 mL) and the combined organic layers were dried over $Na_2SO_4$, filtered, and the solvent was removed in vacuo to yield a colorless solid. The crude material was purified by flash chromatography (0 → 10 % MeOH in $CH_2Cl_2$) to yield oscr#10 (**2**) as a clear colorless film (26.0 mg, 0.085 mmol, 48 % yield). Spectral data for this compound was consistent with that previously reported.[5] TLC (1:9 MeOH:$CH_2Cl_2$ v/v) $R_f$ = 0.30; $^1$H NMR (600 MHz, $CD_3OD$): δ 1.22 (d, *J* = 6.0 Hz, 3H), 1.31-1.44 (m, 8H), 1.55-1.64 (m, 4H), 1.77 (ddd, *J* = 13.5, 11.2, 3.1 Hz, 1H), 1.95 (dt, *J* = 13.1, 3.9 Hz, 1H), 2.27 (app t, *J* = 7.4 Hz, 2H), 3.41 (dt, *J* = 9.5, 6.2 Hz, 1H), 3.48-3.59 (m, 2H), 3.68 (dt, *J* = 9.5, 6.6 Hz, 1H), 3.76 (app s, 1H), 4.49 (s, 1H); $^{13}$C NMR (150 MHz, $CD_3OD$): δ 18.1, 26.3, 27.3, 30.3, 30.37, 30.38, 30.7, 35.4, 36.0, 68.30, 68.34, 69.4, 70.9, 100.4, 178.3; IR (thin film) 3368 (br, m), 2931 (m-s), 2856 (m), 1708 (m), 1048 (s) $cm^{-1}$.

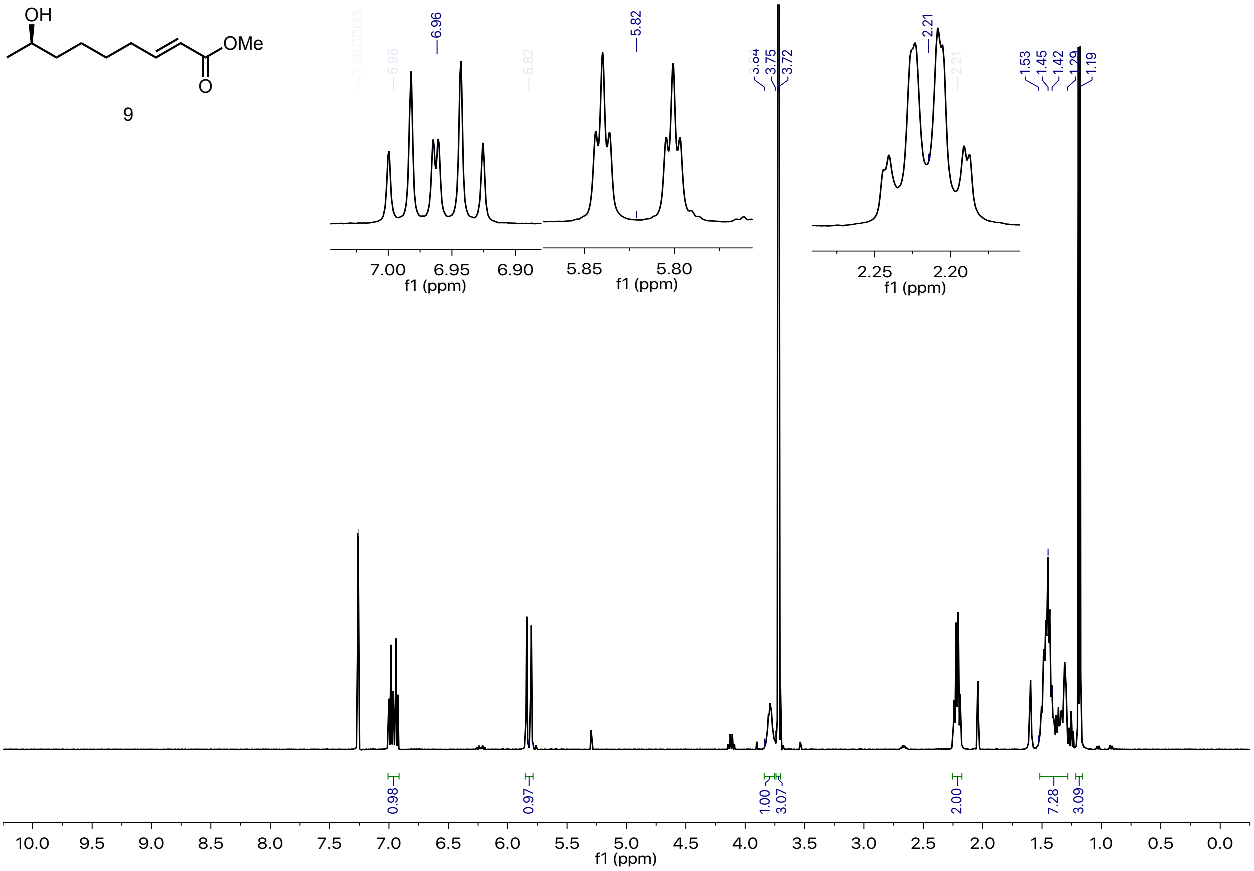


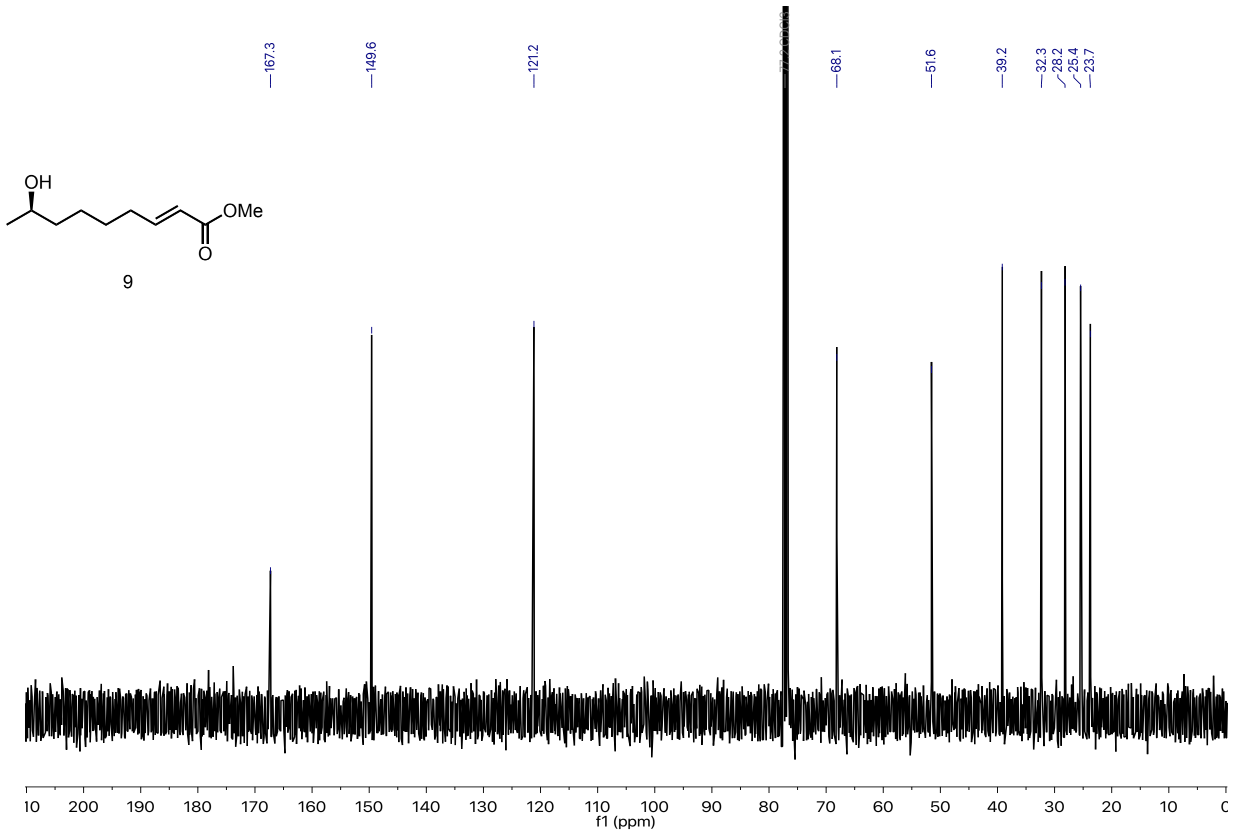


[1]H (400 MHz) and [13]C (100 MHz) NMR of methyl (8*R,2E*)-8-hydroxynon-2-enoate (**9**).

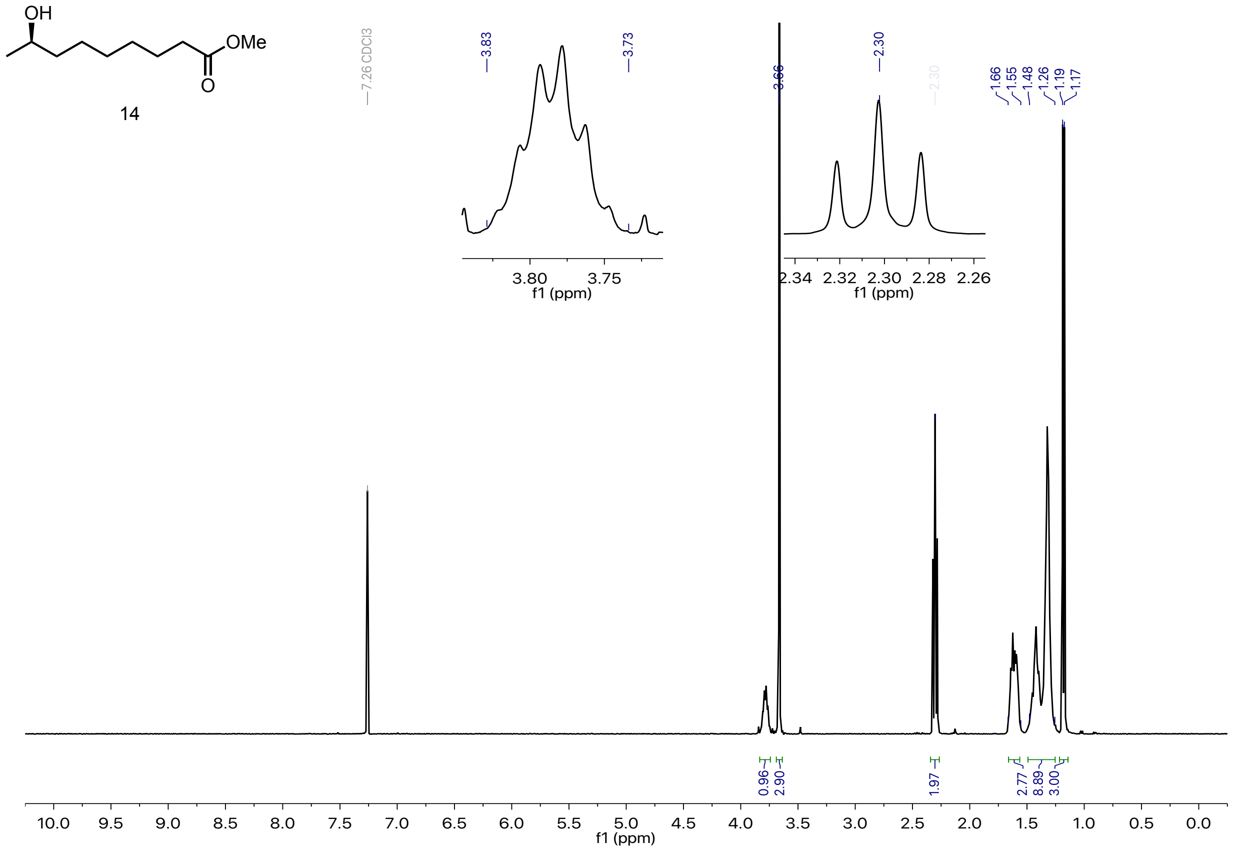


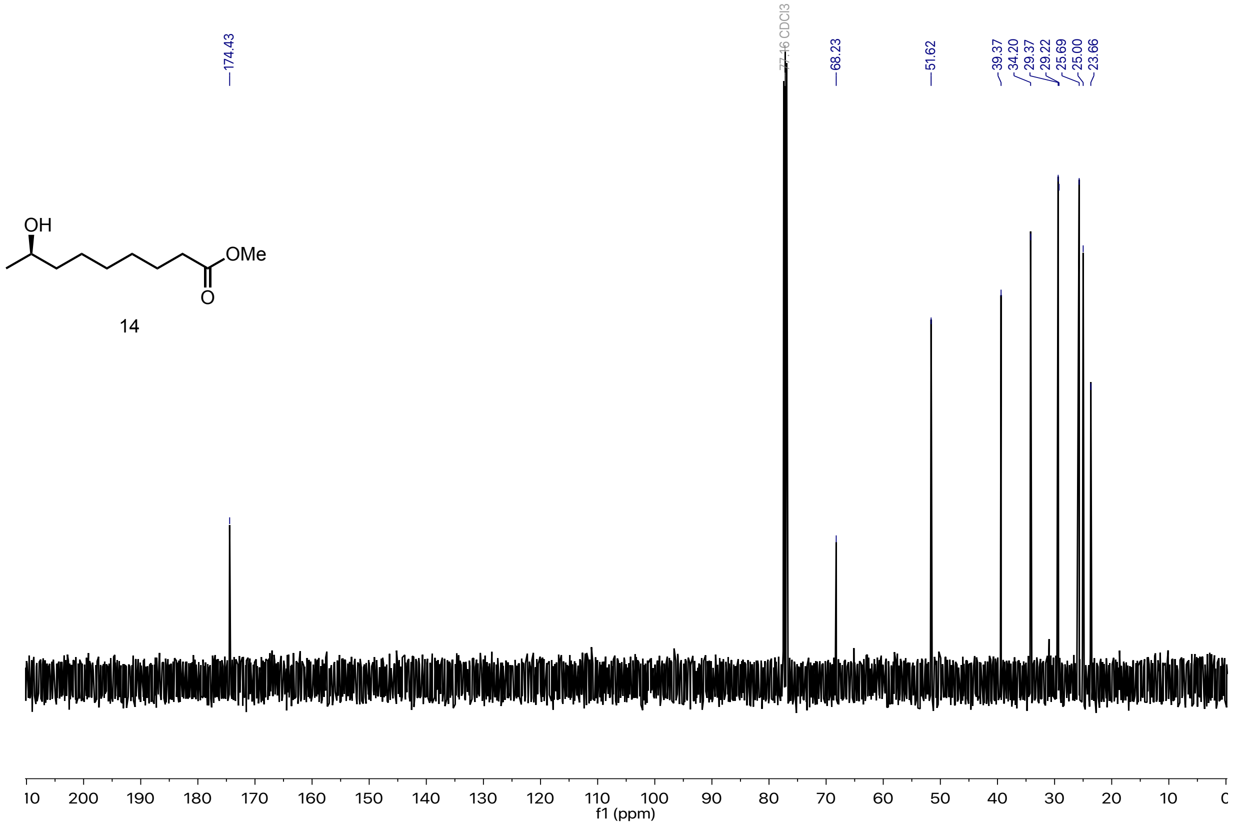


$^{1}H$ (400 MHz) and $^{13}C$ (100 MHz) NMR of methyl (*R*)-8-hydroxynonanoate (**14**).

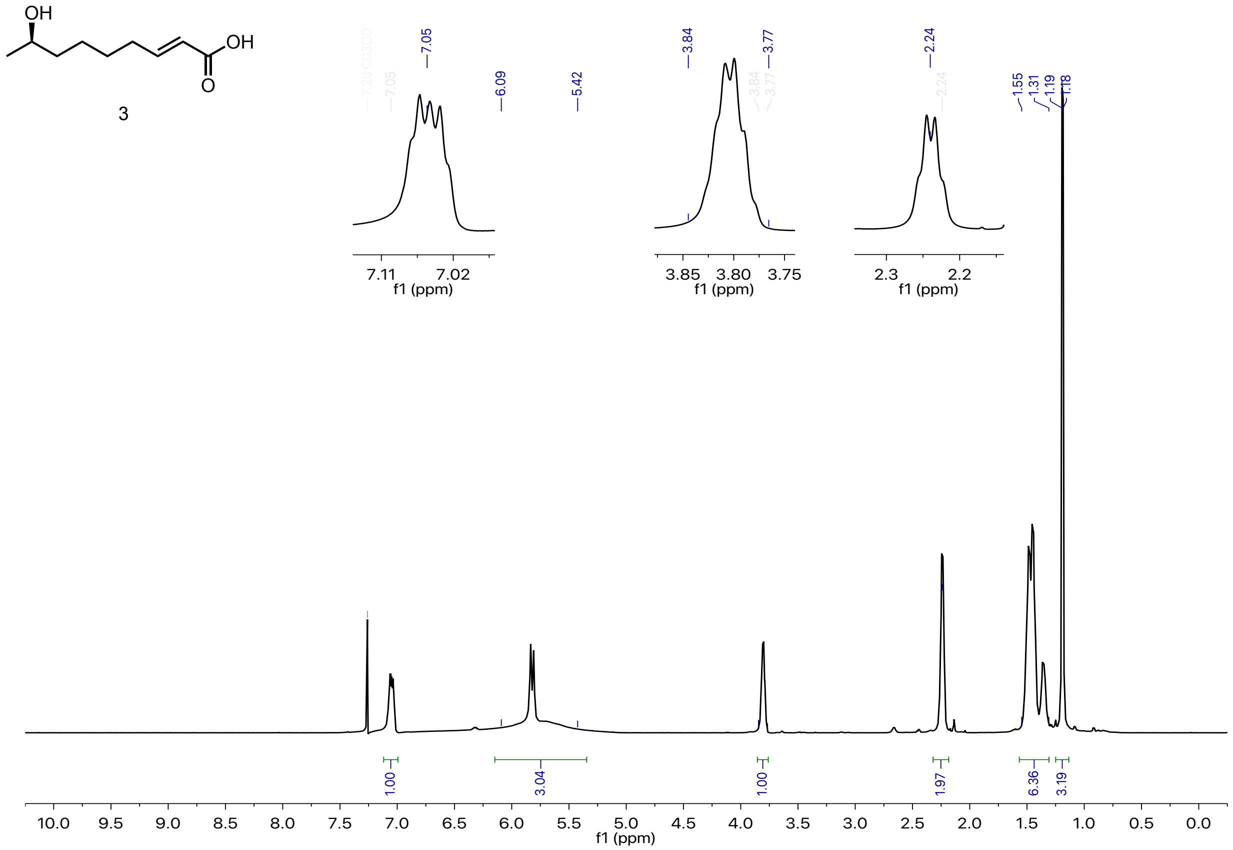

3



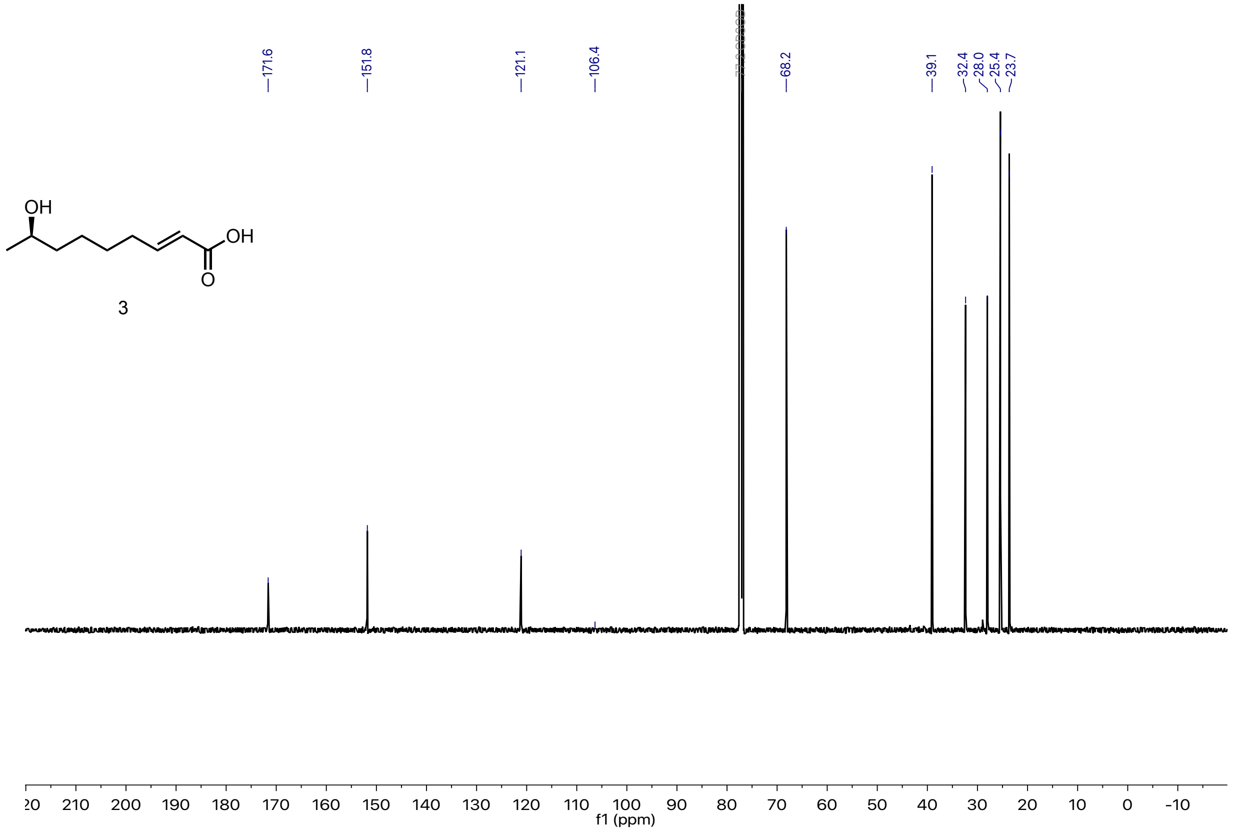

3



$^{1}H$ (600 MHz) and $^{13}C$ (150 MHz) NMR of (8*R*,2*E*)-8-hydroxynon-2-enoic acid (**3**).

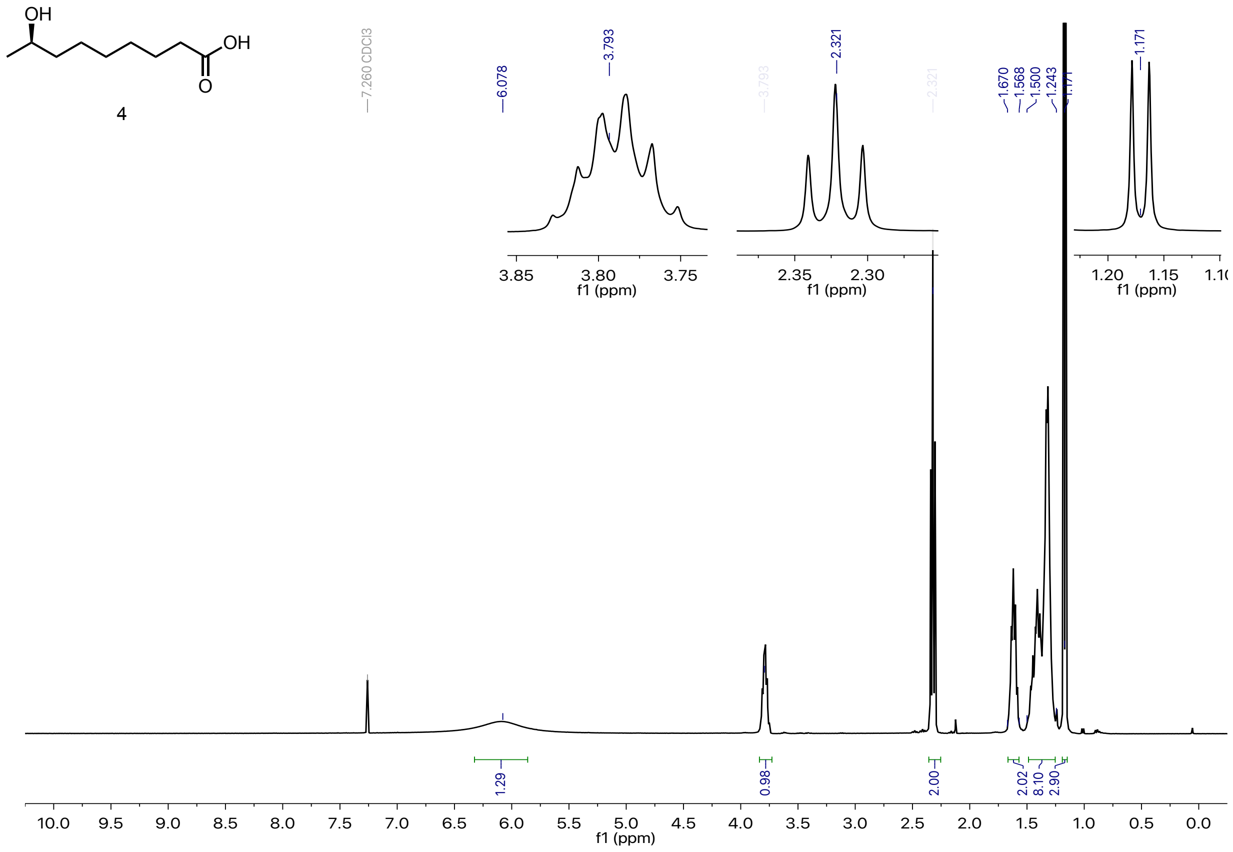


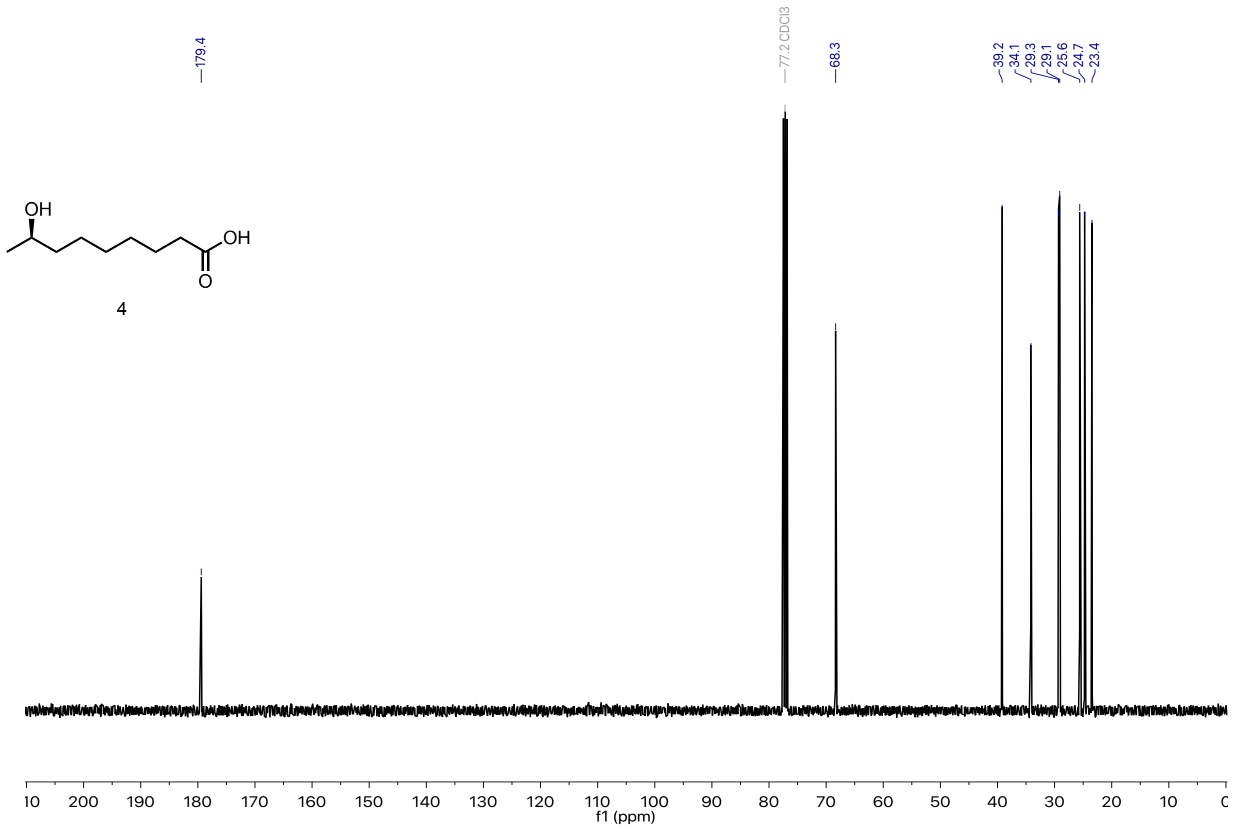


$^{1}H$ (400 MHz) and $^{13}C$ (100 MHz) NMR of (*R*)-8-hydroxynonanoic acid (**4**).

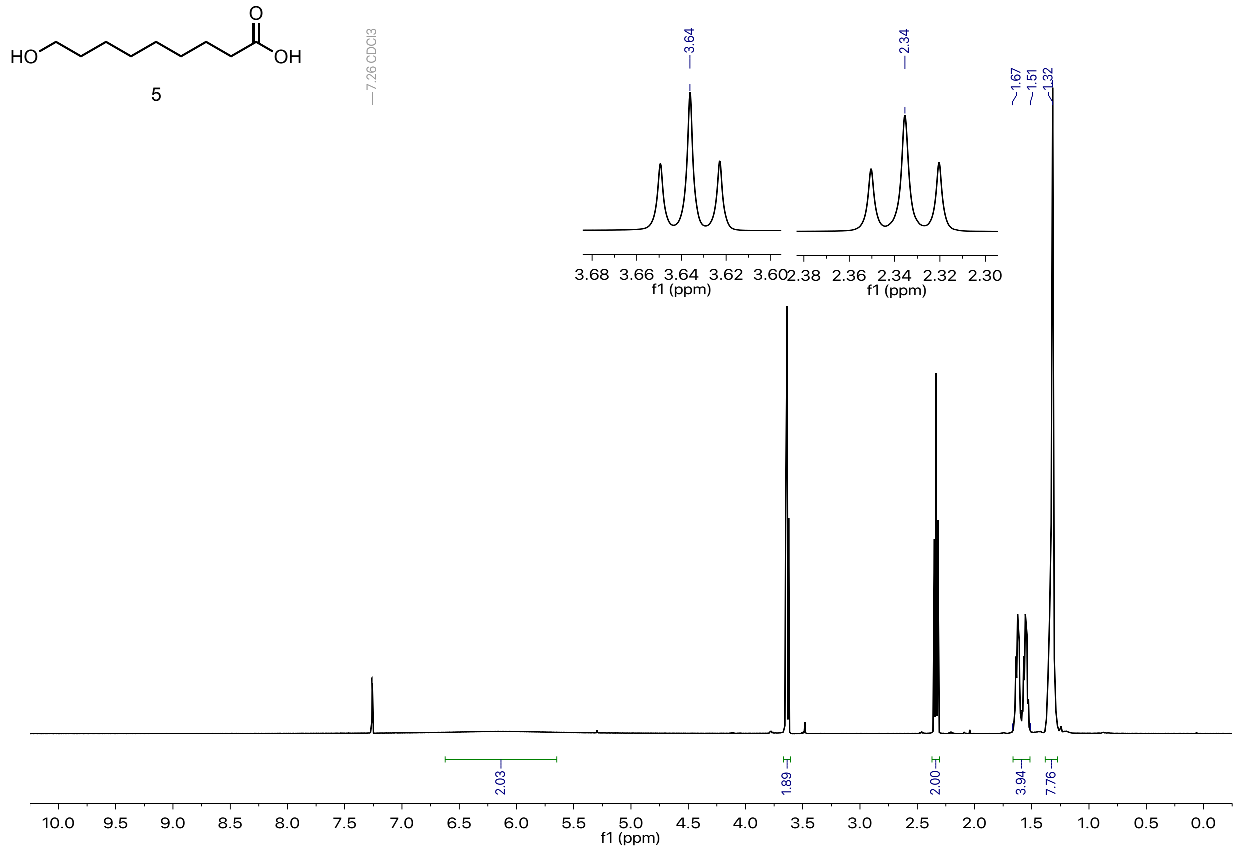


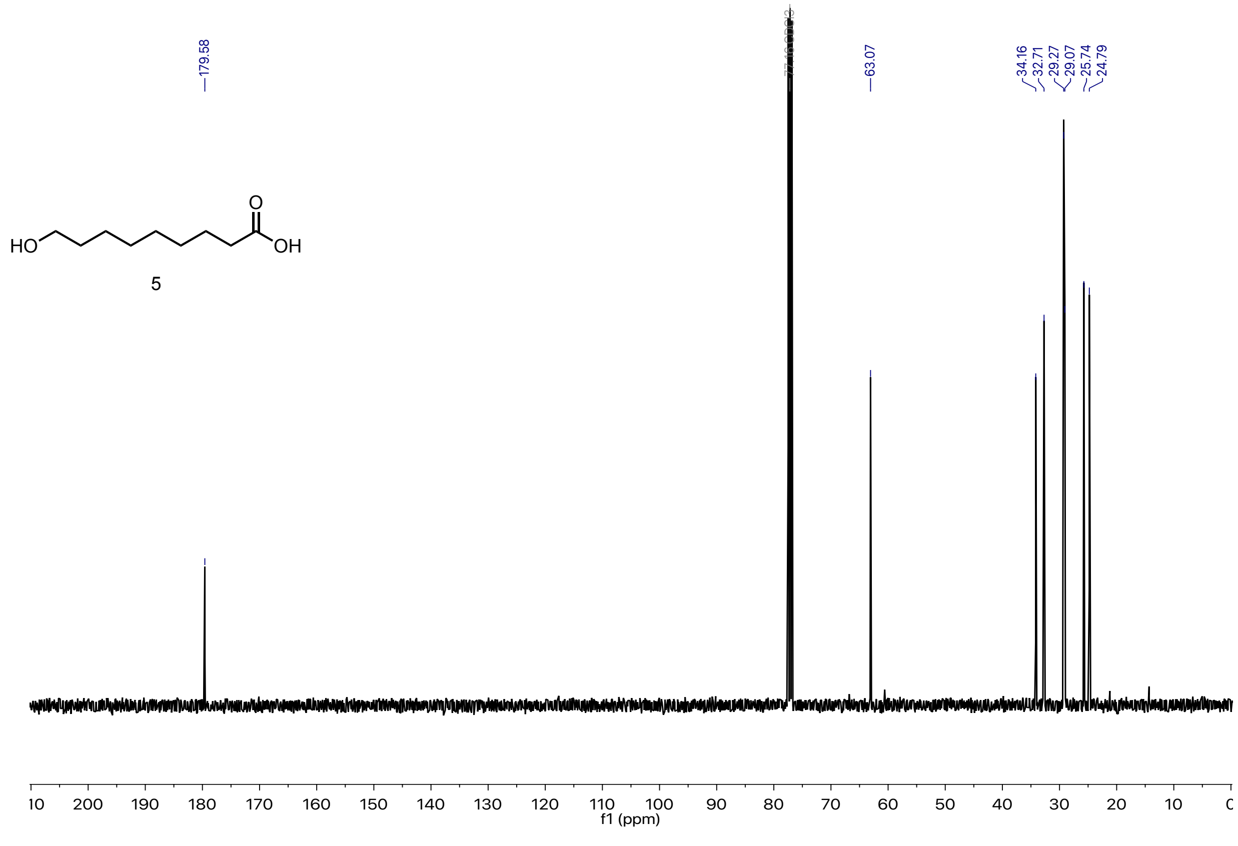


$^{1}H$ (500 MHz) and $^{13}C$ (100 MHz) NMR of 9-hydroxynonanoic acid (**5**).

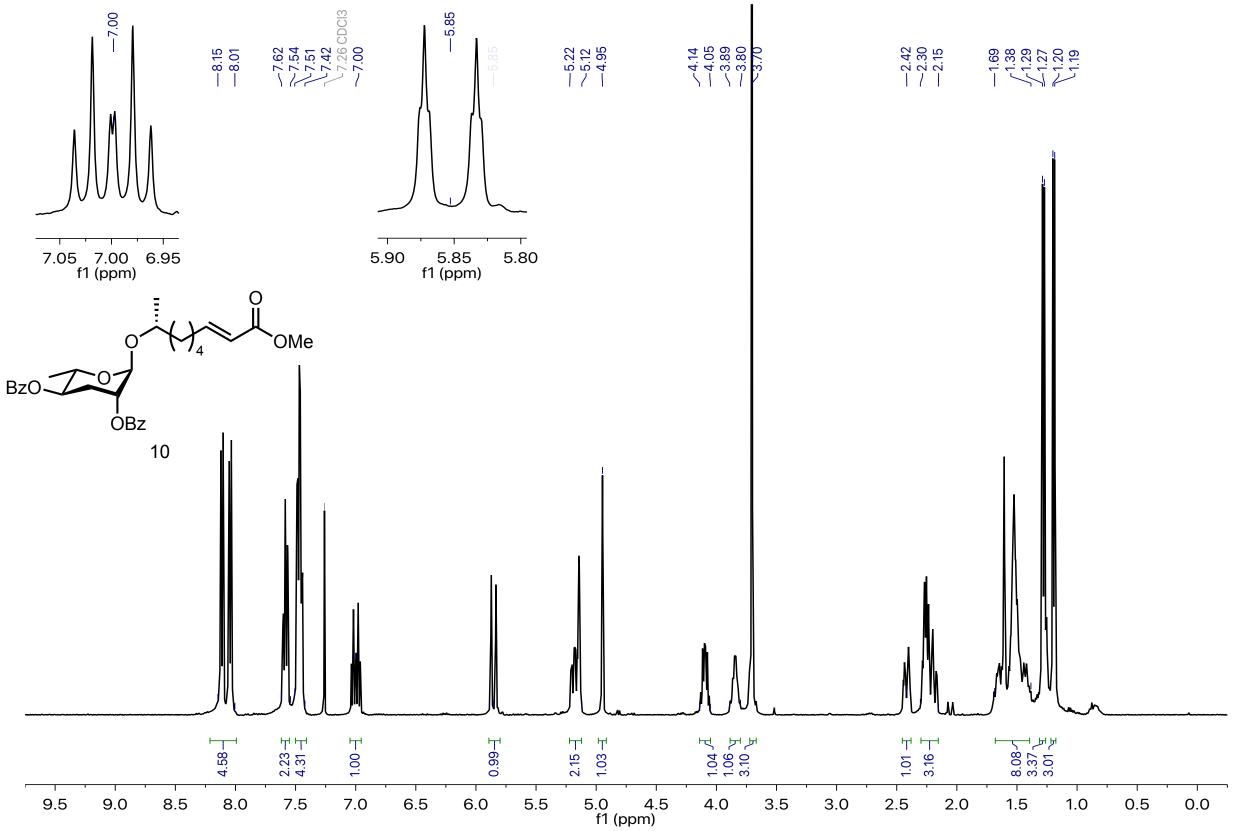


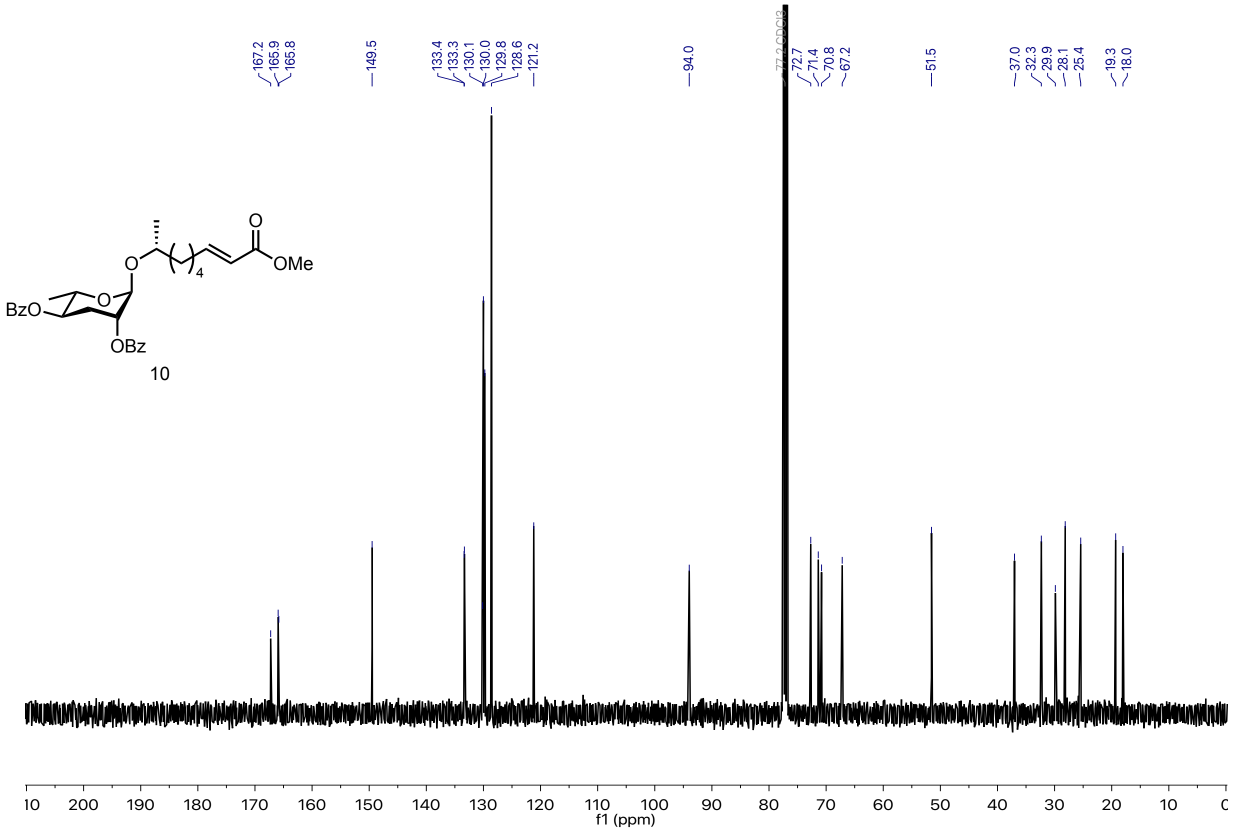


$^{1}H$ (400 MHz) and $^{13}C$ (100 MHz) NMR of methyl ester (**10**).

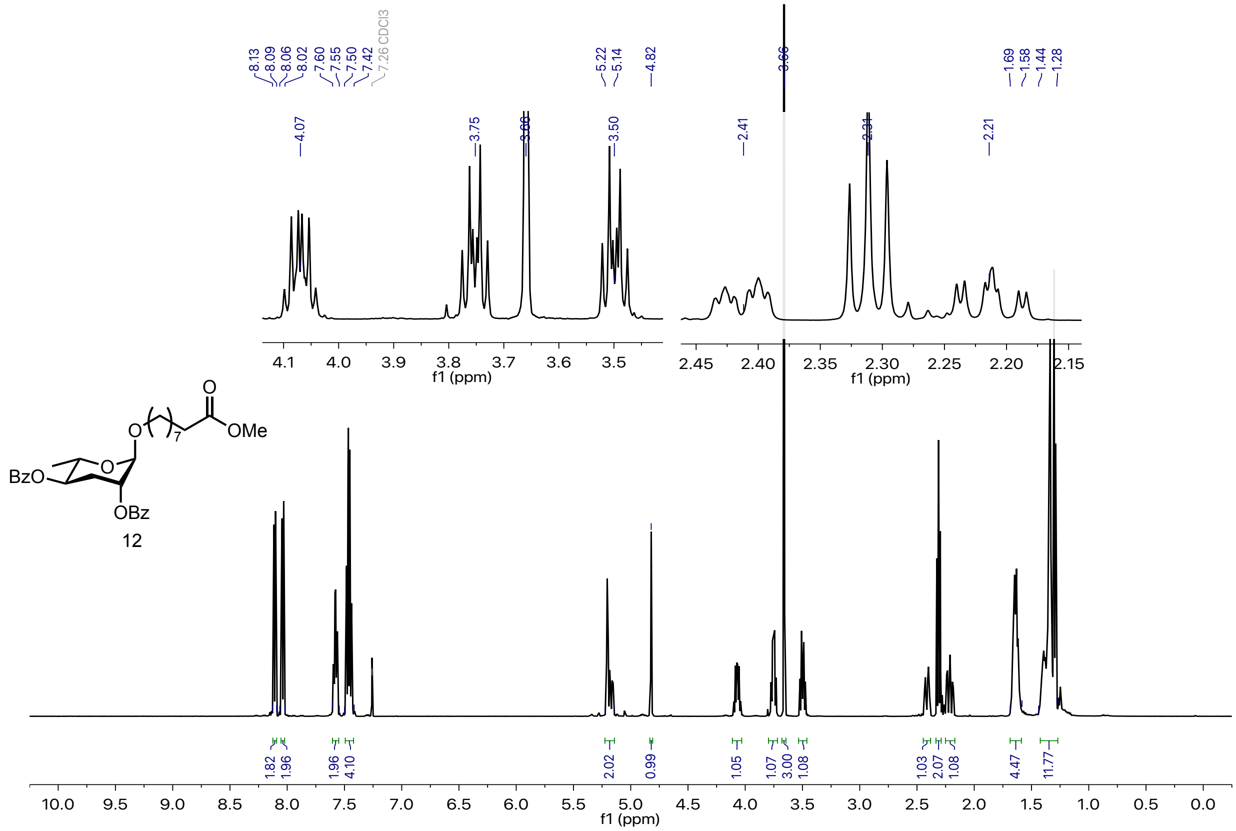


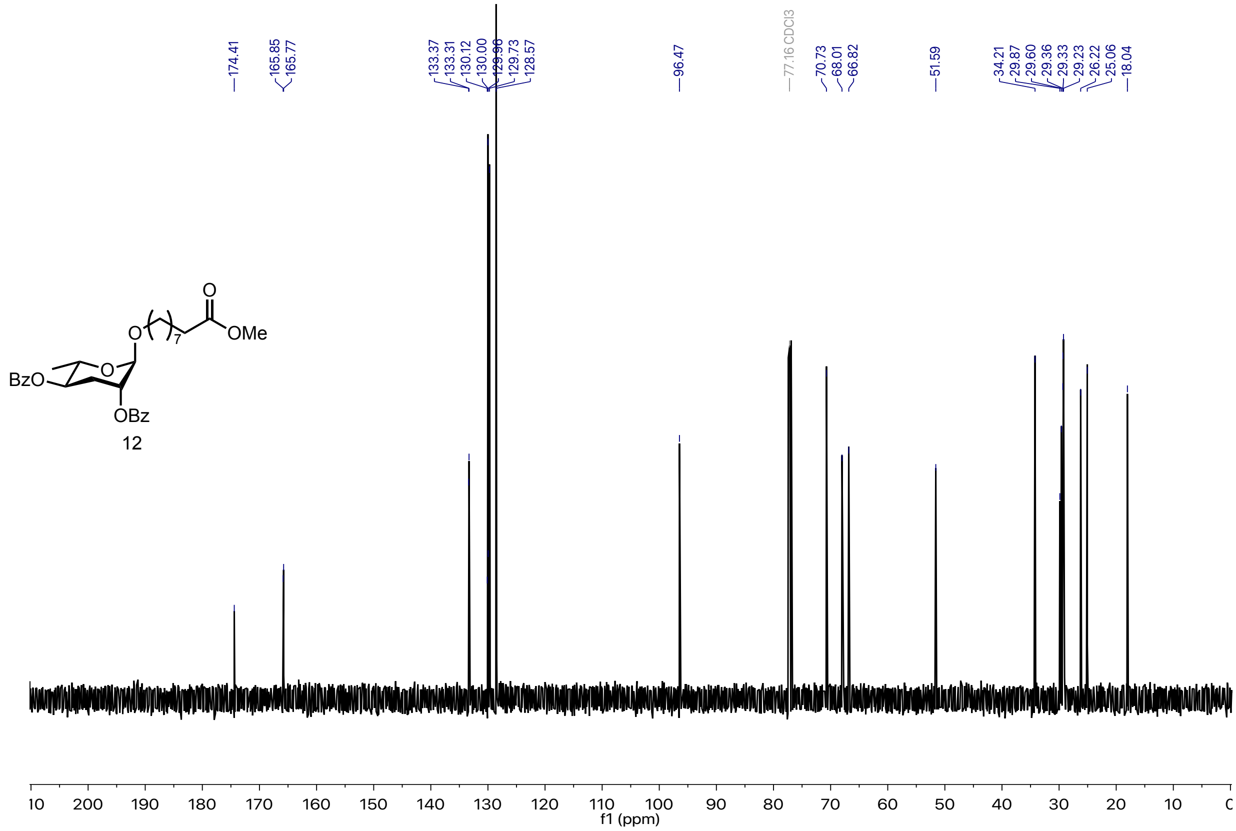


$^1$H (500 MHz) and $^{13}$C (125 MHz) NMR of methyl ester (**12**).

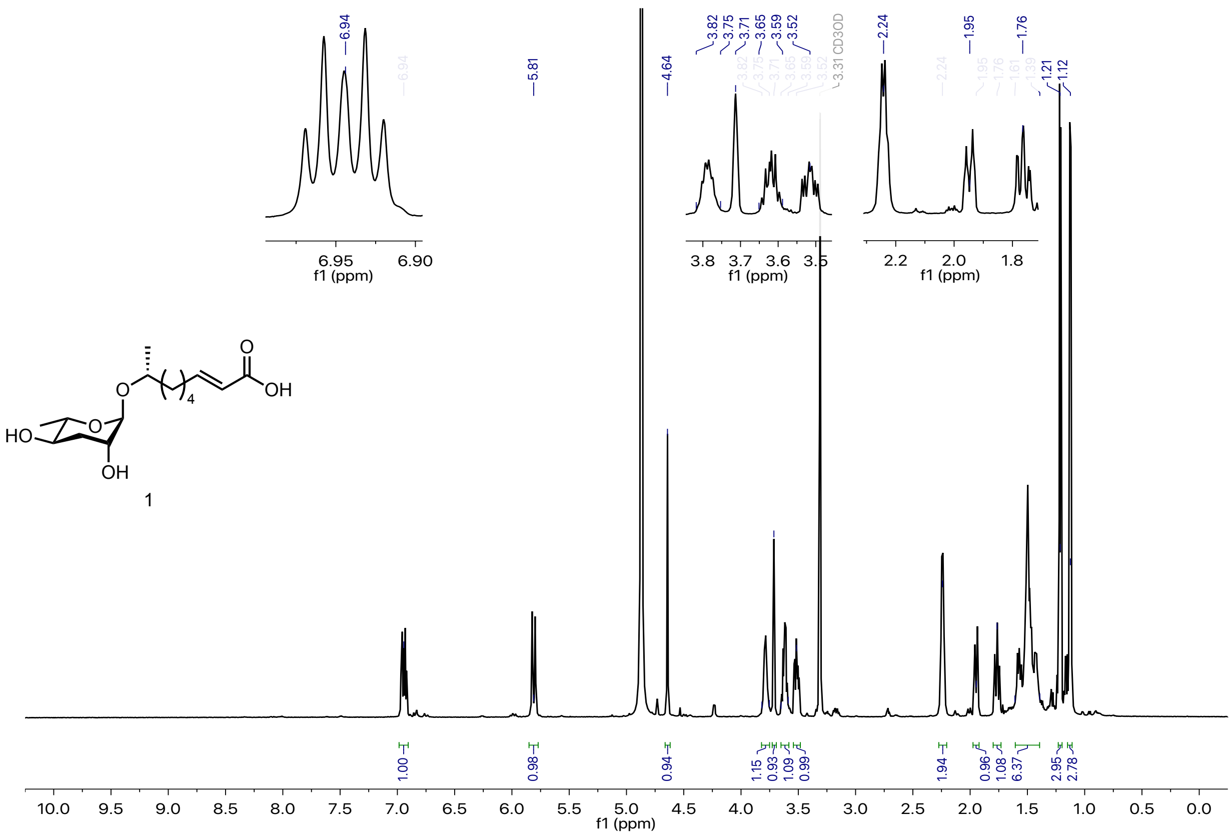


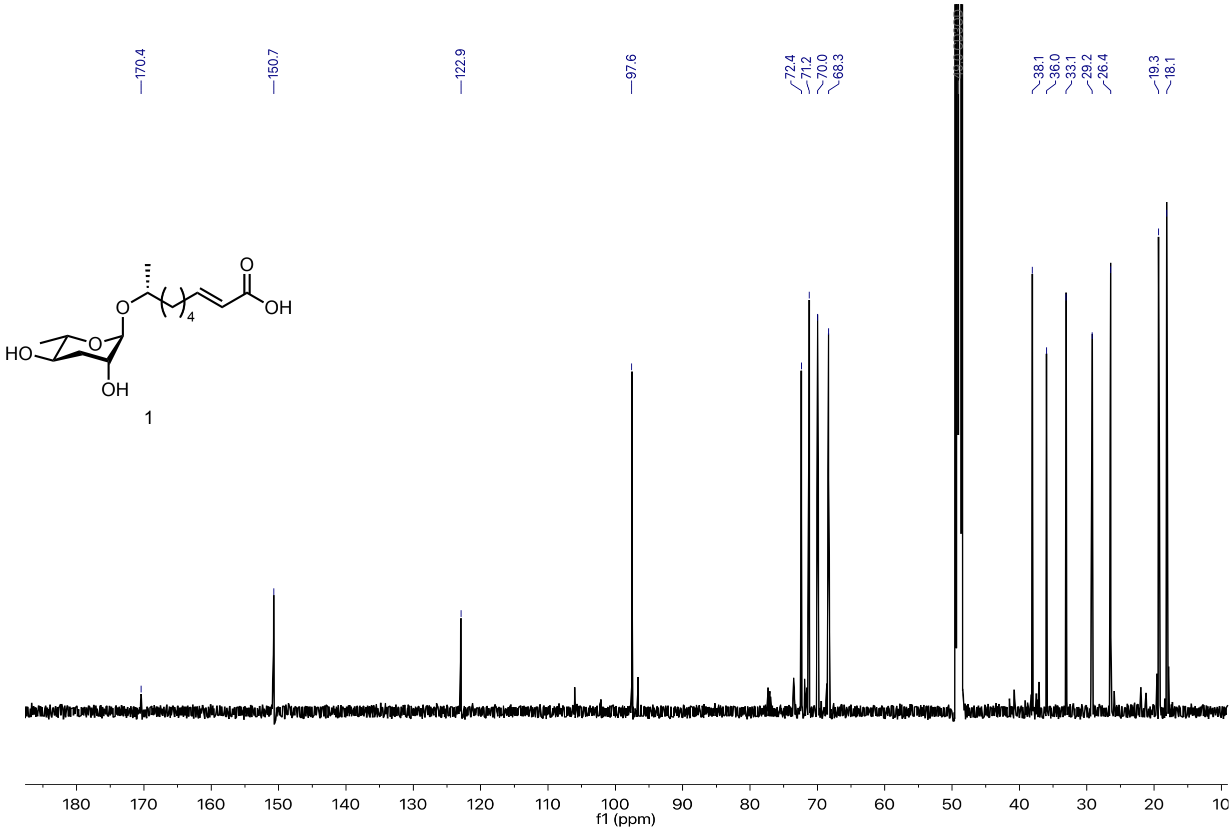


$^{1}$H (600 MHz) and $^{13}$C (150 MHz) NMR of ascr#3 (**1**).

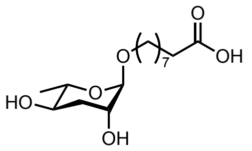


2

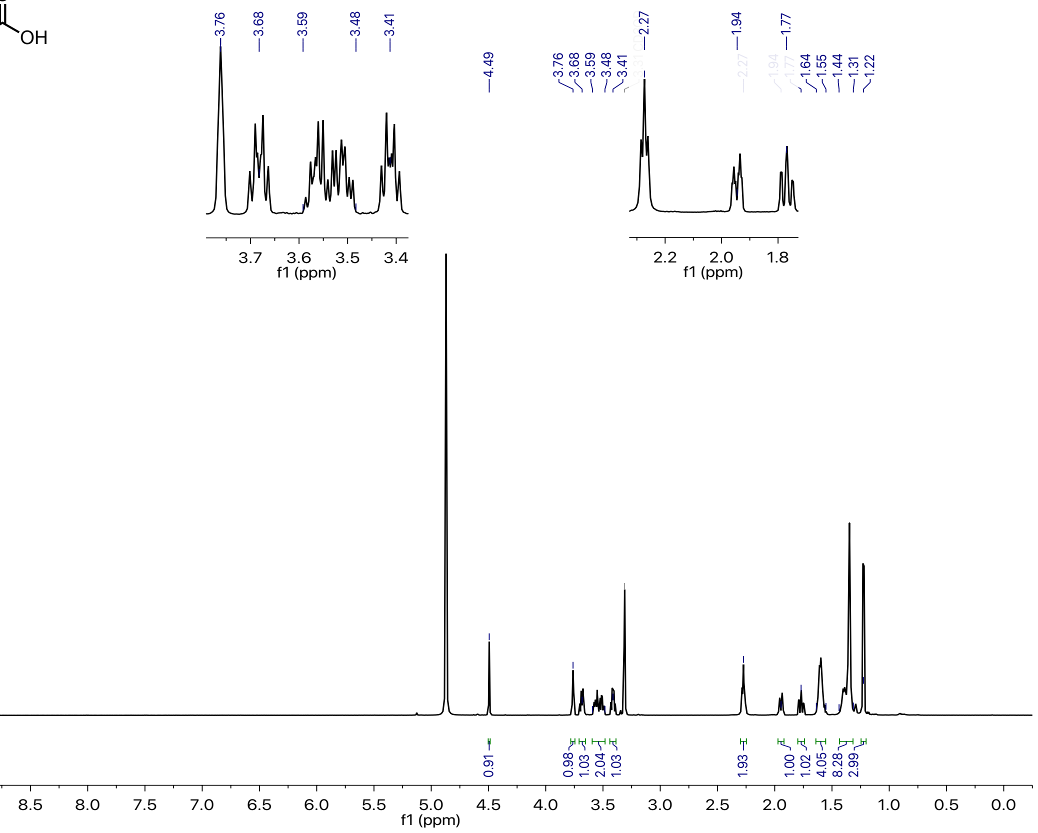


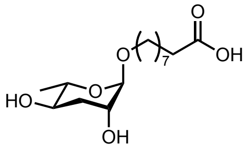


2

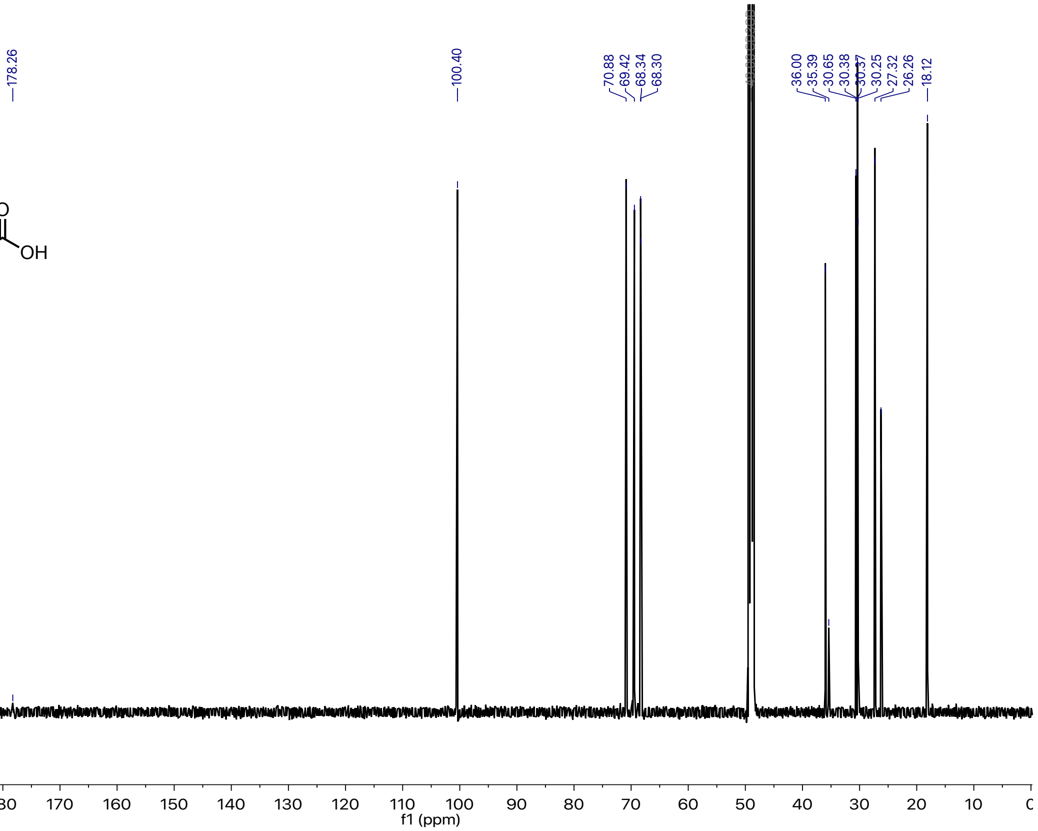


¹H (600 MHz) and ¹³C (150 MHz) NMR of oscr#10 (**2**).
